# *P*-elements strengthen reproductive isolation within the *Drosophila simulans* species complex

**DOI:** 10.1101/2020.08.17.254169

**Authors:** Antonio Serrato-Capuchina, Emmanuel R. R. D’Agostino, David Peede, Baylee Roy, Kristin Isbell, Jeremy Wang, Daniel R. Matute

**Author notes:** Correspondence: Biology Department, University of North Carolina, Chapel Hill, North Carolina, 250 Bell Tower Drive, Genome Sciences Building, Chapel Hill, NC, 27510, USA.

## Abstract

Determining mechanisms that underlie reproductive isolation is key to understanding how species boundaries are maintained in nature. Transposable elements (TEs) are ubiquitous across eukaryotic genomes. However, the role of TEs in modulating the strength of reproductive isolation between species is poorly understood. Several species of *Drosophila* have been found to harbor P-elements (PEs), yet only *D. simulans* is known to be polymorphic for their presence in wild populations. PEs can cause reproductive isolation between PE-containing (P) and PE-lacking (M) lineages of the same species. However, it is unclear whether they also contribute to the magnitude of reproductive isolation between species. Here, we use the *simulans* species complex to assess whether differences in PE status between *D. simulans* and its sister species, which do not harbor PEs, contribute to multiple barriers to gene flow between species. We show that crosses involving a P *D. simulans* father and an M mother from a sister species exhibit lower F1 female fecundity than crosses involving an M *D. simulans* father and an M sister-species mother. Our results suggest that the presence of PEs in a species can strengthen isolation from its sister species, providing evidence that transposable elements can play a role in reproductive isolation and facilitate the process of speciation.

**IMPACT SUMMARY:** Transposable elements (TEs) are repetitive genetic units found across the tree of life. They play a fundamental role on the evolution of each species’ genome. TEs have been implicated in diversification, extinction, and the origin of novelty. However, their potential role in contributing to the maintenance of species boundaries remains largely understudied. Using whole genome sequences, we compared the relative content of TEs across the three species of the *Drosophila simulans* complex. We find that the presence of one TE, *P*-element, in *D. simulans*, and its absence in the sister taxa, differentiates the three species. *P*-elements (PEs) cause a suite of fitness defects in *Drosophila* pure-species individuals if their father has PEs but their mother does not, a phenomenon known as hybrid dysgenesis (HD). We thus studied the possibility that PEs enhance isolation between recently-diverged species. In particular, we studied whether the progeny from interspecific crosses were more prone to suffer from HD than pure species. We found that the presence of paternal PEs reduces hybrid female fecundity, mirroring observations of HD described within species. The effect of PEs is stronger in the interspecific hybrids than in pure species. Our results suggest that PEs can strengthen reproductive isolation in well-formed sister species that still hybridize in nature and pose the question of whether other TEs are involved in the formation of species or in their persistence over time.

## INTRODUCTION

Barriers to gene flow can be categorized depending on where in the reproductive cycle they occur (Dobzhansky 1937; Ramsey *et al*. 2003; Sobel *et al*. 2010). Prezygotic isolation includes all reproductive isolation (RI) traits that preclude the formation of a hybrid zygote, such as ecological, behavioral, and gametic incompatibilities (Dobzhansky, 1937). Postzygotic barriers occur after a hybrid zygote is formed and include phenotypes as extreme as hybrid sterility and inviability (Orr and Presgraves 2000; Orr 2005), but also more nuanced traits, such as hybrid behavioral defects (McQuillan *et al*., 2018; Turissini *et al*., 2017) and delays in development (Matute and Coyne 2010; Ishikawa *et al*. 2011). While both types of barriers can lead to speciation, prezygotic barriers tend to be completed faster than postzygotic ones (Sasa *et al*. 1998; Moyle *et al*. 2004; Turissini *et al*. 2018). Nonetheless, postzygotic barriers seem to be crucial to the persistence of species in the face of gene flow once lineages come into secondary contact (Rosenblum *et al*., 2012; Coughlan and Matute 2020).

The genetic basis of postzygotic isolation usually stems from deleterious interactions between parental genomes in hybrids, which can occur at the allelic level (reviewed in (Maheshwari and Barbash, 2011; Nosil and Schluter, 2011)) or the genomic level (i.e., ‘genome clashes’; (Nolte *et al*. 2009; Dion-Côté *et al*. 2014; Dion-Côté and Barbash 2017)). Transposable elements (TEs) are a candidate to cause the latter (Ginzburg *et al*. 1984; Castillo and Moyle 2012; Watson and Demuth 2013; Serrato-Capuchina and Matute 2018). TEs are repetitive genetic units found ubiquitously across taxa and have been linked to molecular and phenotypic novelty (McClintock 1953; Werren 2011). TEs can facilitate rapid genotypic change by causing genic interruptions, duplications, or gene expression changes through their transposition (Trizzino *et al*. 2017). In pure species, TEs are repressed by the interaction between PIWI genes and non-coding RNAs called Piwi-interacting RNAs (piRNAs). piRNAs are small (24–35 nucleotides long) and suppress TE expression across multiple taxa (reviewed in Michalak, 2009). Derepression of TEs through a mismatch of regulatory systems might activate dormant transposons, which in turn can have deleterious fitness implications in hybrids. Comparative gene expression surveys suggest TE misexpression is common in hybrids but not in pure species (Pélissier *et al*. 1995, 1996; Metcalfe *et al*. 2007; Dion-Côté *et al*. 2014), which might be an indication of the role of TE derepression in hybrid breakdown.

*P*-elements (PEs) in *Drosophila melanogaster* is one of the best-studied TEs in terms of molecular action and phenotypic consequences (reviewed in (Kelleher 2016)). The invasion of PEs in all populations of *D. melanogaster* occurred in the past 40 years (Kidwell *et al*. 1977). PEs in *D. melanogaster* cause a variety of phenotypes collectively named hybrid dysgenesis (HD), a syndrome of deleterious effects in F1s from crosses between a mother who lacks PEs (♀M) and a father who carries them (♂P). F1s from these crosses show a suite of defects that include sterility, chromosomal breaks, and increased mutation rates (Kidwell and Kidwell 1975; Kidwell *et al*. 1977; Schaefer *et al*. 1979; Engels and Preston 1980; Kidwell 1983a). Conversely, in crosses where the mother carries PEs (♀P), F1s do not show hybrid dysgenesis, regardless of the genotype of the father, since piRNAs and associated Piwi proteins inherited from the mother confer protection from deleterious effects of paternally-inherited PEs. This interaction between PEs and maternally-inherited piRNAs ultimately determines whether an F1 suffers from hybrid dysgenesis due to this TE (Slotkin and Martienssen 2007; Siomi *et al*. 2011; Saito 2013).

While it is clear that PEs can cause hybrid dysgenesis within *D. melanogaster* and *D. simulans* (Kidwell *et al*. 1977; Bingham *et al*. 1982; Hill *et al*. 2016), the role of PEs in contributing to RI between well-formed species remains largely untested. If two potentially hybridizing species differ in their TE content (e.g., PEs), hybrid progeny might suffer from HD. Examining the potential effect of the presence vs. absence of PEs on RI between species requires the study of a group of species in which: *i*) some species harbor PEs while others do not, *ii*) species hybridize, and *iii*) produce fertile progeny. *Drosophila melanogaster*, the classical model for the study of HD, can intercross with several other species, but all viable hybrids are sterile and consequently does not fulfill these criteria (but see (Davis *et al*. 1996; Barbash and Ashburner 2003; Brideau *et al*. 2006)). Species from the *willistoni* and *saltans* species groups seem to all have PEs (Mizrokhi and Mazo 1990; Clark *et al*. 1995; Brookfield and Badge 1997; Silva and Kidwell 2000; De Setta *et al*. 2007) and thus do not satisfy the criteria either.

The *simulans* species group, which contains *D. simulans, D. sechellia* and *D. mauritiana*, is an ideal system to assess the potential effects of TEs in RI for at least three reasons. First, PEs recently invaded *D. simulans* (Kofler *et al*. 2015) and are only present in some lines. While most *D. simulans* individuals collected after 2015 harbor functional PEs, neither *D. sechellia* nor *D. mauritiana* shows any evidence for their presence, past or present (Kofler *et al*. 2015). Second, all pairwise crosses of species in this group produce fertile female F1 hybrids (Lachaise *et al*. 2018; Turissini *et al*. 2018). Third, there is evidence of hybridization and admixture in nature among these three species. *Drosophila simulans* and *D. sechellia* hybridize in the central islands of the Seychelles archipelago (Matute and Ayroles 2014) and show signatures of gene exchange (Garrigan *et al*. 2012; Schrider *et al*. 2018). Despite the absence of a contemporary hybrid zone between *D. mauritiana* and *D. simulans*, these two species also show evidence of introgression (Garrigan *et al*. 2012; Brand *et al*. 2013; Meiklejohn *et al*. 2018). The *simulans* species group is thus a powerful system to assess whether TEs modulate the strength of RI in nature because the natural variation in *D. simulans* lines allows for the test of whether populations with PEs show stronger RI towards the sister species than populations without PEs.

Here, we leverage the recent PE invasion of *D. simulans* to test whether the presence of PEs affects the magnitude of postzygotic reproductive isolation between species that differ in their PE content. We first compare relative content of TEs across species to assess the extent to which PE uniquely explains TE variance across species. Next, we find that PEs do affect F1 female fecundity, mirroring observations of hybrid dysgenesis described within species (i.e., hybrid dysgenesis only occurs in ♀M/♂P F1s). This effect is only observable in F1 females, as F1 males from interspecific crosses between species of the *simulans* species complex are invariably sterile. As observed in pure species, progeny from fathers with higher PE copy numbers show stronger HD, but this effect is stronger in the interspecific hybrids. Our results serve as a formal test for the putative role of PEs in strengthening RI between sister species.

## MATERIALS AND METHODS

### TE detection: sequencing and read mapping

i. **Genome sequencing, short reads**: For assessing the number of TE copies in each of the three species in the *simulans* complex, we used short reads of 115 lines (42 *D. sechellia*, 15 *D. mauritiana*, and 58 *D. simulans*). Table S1 describes which lines were previously sequenced and lists the accession numbers of all lines. To sequence new lines (two *D. sechellia* and two *D. mauritiana*), we obtained genomic DNA using the Qiagen^®^DNeasy^®^ 96 Tissue Kit following the protocol and recommendations from the manufacturer. Next, we outsourced library construction and sequencing to the High Throughput Sequencing Facility at the University of North Carolina, Chapel Hill. The libraries were built using the Nextera protocol as specified by the manufacturer. We obtained read quality information using HiSeq Control Software 2.0.5 and RTA 1.17.20.0 (real-time analysis). CASAVA-1.8.2 generated and reported run statistics of each of the final FASTQ files. Resulting reads ranged from 100bp to 150bp, and the target average coverage for each line was 20X.
ii. **Genome sequencing, single-molecule sequencing:** We sequenced two *D. sechellia* (Anro71 and Denis72) and two *D. mauritiana* lines (R42 and R50) with Oxford Nanopore Technology (ONT) to assess if any of these lines harbored complete PE copies (see immediately above). For each of these four lines, we extracted DNA from 20 whole male flies of each line using a modified phenol:chloroform protocol (Sambrook and Russell 2006). We prepared libraries using the Ligation Sequencing Kit 1D with native barcoding (SQK-LSK109 and EXP-NBD104, Oxford Nanopore) according to the manufacturer’s protocol. Libraries were sequenced on six R9.4 flow cells with an average of 1,526 available pores and run for 48 hours, or until no pores were available, on a GridION running MinKNOW v3.1.8. Reads were basecalled using Guppy (ONT) with the dna_r9.4.1_450bps_flipflop model. For each line, we mapped reads in a pairwise fashion using Minimap2 v2.15-r905 (Li 2018) and assembled them using Miniasm v0.3-r179 (Li 2016). We then corrected and generated consensus assemblies with four iterations of Racon v1.3.2 (Vaser *et al*., 2017), followed by Medaka v0.6.2 (https://github.com/nanoporetech/medaka).
iii. **TE relative frequency.** We aligned Illumina short-read data from the lines sequenced to a list of canonical *Drosophila* transposable element sequences (v10.1, https://github.com/bergmanlab/transposons/blob/master/current/transposon_sequence_set.fa) using minimap2 (Li 2018) with the short-read option (“-cx sr”). We then tabulated the number of base pairs from each sample aligning to each transposable element and calculated the effective copy number per TE per sample by first dividing that quantity by the canonical length of the TE. This, in turn, was normalized for the depth of sequencing by dividing by the ratio of total base pairs sequenced to genome size. For the latter, we used a fixed estimate of 125 Mb. We generated consensus sequences per TE per line, defining each base as the major allele mapping to that site and leaving as undefined any site for which no allele had more than five supporting reads. This approach differs from previous approaches (Serrato-Capuchina *et al*. 2020), in that we used the total number of base pairs to calculate the TE copy number and not the number of mapped reads. The results are similar (see below). If a line showed fewer than 0.5 copies of the PEs (i.e., a single heterozygote PE copy), it was considered PE-negative. To assess whether our inference was affected by the way we scored copy number, we tried two different cutoff schemes and counted a copy as complete if it was missing (i.e., the alignment to the reference contained ambiguous bases for) no more than than 0% and 25% of the sequence. This approach potentially reveals the number of copies in each isofemale at the time of sequencing. We verified the results from this approach with the long-read sequencing.

### TE detection: statistical analyses

To determine which TEs differ in their content across the three species of the *simulans* species complex, we used a linear discriminant analysis (LDA) for the species group. We used the function ‘*lda*’ (library *MASS*, (Venables *et al*., 2003)) to find the best two linear discriminant (LD) functions, LD1 and LD2. For these analyses, we only retained TEs for which we had inferred at least one full copy in more than each sequenced line. We plotted the scores of these two functions using the function *plot* (library *graphics*, (R Core Team 2016)). We extracted the relative contribution of each PE from the object *scaling*. To determine which TEs differed between species, we fit nested ANOVAs for each TE using *lm* (library ‘*stats*’, (R Core Team 2016)). For all these linear models, estimated copy number was the response, species was a fixed factor, and the line was a nested effect within species. We then performed pairwise comparisons using a Tukey HSD test using the function *glht* (library ‘*multcomp*’, (Hothorn *et al*., 2008)). We present analyses for the PE copy number estimation using mapped reads (Serrato-Capuchina *et al*. 2020) and total mapped based pairs (described immediately above).

### Fly stocks

All experiments reported in this manuscript used isofemale lines; all the lines have been previously used and reported elsewhere. We used at least seven lines for each species of the *simulans* species group. The details of each line are listed in Table S2 and are described by species as follows.

#### D. simulans

PEs have been present in *D. simulans* for the last 15 years and have rapidly increased in frequency during that time (Kofler *et al*. 2015). For crosses, we used seven lines which showed evidence of P-elements (denoted as P): 18MPALA11, cascade1, 18MU05, 18KARI10, 18MPALA13, 18MPALA02C, and BiokoLB1. We also used seven lines with no evidence for the presence of PEs (denoted as M): Md06, Md105, Md106, Md199, NS05, NS40, and NS113. Donors and collections details of the lines are listed in Table S2. These lines were collected in Africa between 2000 and 2019 (Rogers *et al*. 2014; Comeault *et al*. 2017) and vary in their PE load (Serrato-Capuchina *et al*., 2020). All of these lines have been sequenced; Table S1 lists the SRA number of each line.

#### D. sechellia

Unlike *D. simulans, D. sechellia* lacks functional PEs ((Daniels *et al*. 1990; Kofler *et al*. 2015; Hill *et al*. 2016), see results). For crosses (see below), we used seven lines collected on the islands of La Digué, Denis, and Mahé (Anse Royale Beach) by DRM and J.F. Ayroles in 2012: LD11, LD16, Denis72, DenisNF13, DenisNF100, Denis124, and Anro71. Instead of banana-yeast traps, females were collected using bottles seeded with ripe *Morinda* fruits, due to resource preference. Additional details for the lines have been published elsewhere (Matute and Ayroles 2014). Six lines have been previously sequenced (Schrider *et al*., 2018; Turissini *et al*., 2017). We obtained short-read sequencing data for Anro71 and long-read sequencing data for Anro71 and Denis72 (see above). Table S2 lists the collection location, collectors, and SRA number of each line.

#### D. mauritiana

Similar to *D. sechellia, D. mauritiana* shows no evidence of PE presence (Brookfield *et al*. 1984; Daniels *et al*. 1990; Kofler *et al*. 2015). For crosses, we used seven lines collected in the islands of Mauritius and Rodrigues: R8, R23, R31, R32, R61, R42, and R50. Donors and details of the lines are listed in Table S2. The first five lines have been previously sequenced (Brand *et al*., 2013; Garrigan *et al*., 2012; Matute *et al*., 2020; Meiklejohn *et al*., 2018). We obtained sequencing data (short- and long-read) for R42 and R50 (see above).

### F1 Production

Our goal was to study the effect of TEs in interspecific F1 fecundity (see below) at two different temperatures. In all instances, we present all crosses ordered as female × male. For ♀M × ♂P and ♀M × ♂M crosses, we crossed females from seven lines of each of M *D. simulans*, seven *D. mauritiana*, and seven *D. sechellia* lines to males from seven M and seven P *D. simulans* lines for a total of 294 types of crosses (21 × 14). For ♀P × ♂M crosses, we crossed females from seven lines of each of M and P *D. simulans* to seven lines of each of M *D. simulans, D. sechellia*, and *D. mauritiana*, for a total of 294 types of crosses (14 × 21). For both within- and between-species crosses, we placed four to nine day-old virgins of each species in 30mL bottles with yeast and cornmeal medium. For conspecific crosses, we used a 1:1 ratio of females to males in each cross. For heterospecific crosses, we used a 1:2 ratio of females to males in each cross to account for greater female choosiness (Lachaise *et al,*. 1986; Turissini *et al*., 2018), with no more than 50 total flies per vial. Right after mixing, we maintained the vials at 24°C for 24 hours. After this time, vials were placed in a 25°C or 29°C incubator (Percival DR-36VL; Percival Scientific Inc.; Boone, USA). Every six days, we transferred the adults to a new vial and inserted a pupation substrate (KimWipes, Kimberly-Clark; Roswell, USA) treated with 0.5% propionic acid to the old vial. Vials were inspected daily for the production of progeny. F1s produced were collected for up to 15 days after parentals were removed. Our goal was to study at least 20 individuals per cross. We dissected 20 females for each cross with the exception of the crosses listed in Table S3, for which we dissected 40 females per cross.

### Ovary number, counts

One of the most pronounced phenotypes of the HD syndrome is reduced female fecundity, as evidenced by the presence of atrophied gonads (ovaries). We scored the number of functional ovaries in pure species and F1 hybrids in the *simulans* species complex at two temperatures. The procedure for conspecific and heterospecific crosses was identical. We produced F1 females as described above (See ‘F1 Production’). To score female fecundity, we counted the number of functional gonads in each F1 female (i.e., 0, 1, or 2 ovaries). We collected females aged four to nine days post-eclosion, anesthetized them with CO_2_, and extracted their reproductive tracts using forceps (Wong and Schedl 2006). The ovaries from each individual were then fixed on a pre-cleaned glass slide with chilled *Drosophila* Ringer’s solution (Cold Spring Harbor Protocols, Serrato-Capuchina *et al*. 2020). We counted the number of functional and atrophied gonads for each individual. An ovary was considered functional if it contained at least one ovariole (egg chamber). We used a Leica S6E stereoscopic microscope (Leica Microsystems; Denmark) to score all dissections. We scored 20 females for each of the conspecific and heterospecific combinations (Table S3 lists the exceptions).

### Ovary number, statistical analysis

A distinct trait of HD is a reduced number of ovaries in females from ♀M × ♂P crosses (Kidwell *et al*., 1977; Engels and Preston 1980; Rubin *et al*., 1982; Kidwell 1985; Hill *et al*., 2016). Our goal was to assess whether the presence of PEs led to a more pronounced HD phenotype in heterospecifics than in conspecifics. First, we compared whether different crosses (e.g., ♀ M × ♂ P vs. ♀ M × ♂ M) showed differences in the proportion of dysgenic females they produced. To display the results, we pooled the observations for all crosses that involved the same maternal line and the same type of male (i.e., same species, either ♂M or ♂P), and calculated the binomial confidence interval of each proportion using the function ‘*binom.cloglog*’ (library *binom*, (Dorai-Raj 2015)) and plotted the means and confidence intervals using the function ‘*plotCI*’ (library *gplots*, (Warnes 2012; Warnes *et al*. 2016)). We compared whether proportions of dysgenic progeny differed between ♀*sim*M × ♂*mau* and ♀*sim*P × ♂*mau*, between ♀*sim*M × ♂*sech* and ♀*sim*P × ♂*sech*, between ♀*mau* × ♂*sim*M and ♀*mau* × ♂*sim*P, and between ♀*sech* × ♂*sim*M and ♀*sech* × ♂*sim*P using two-sample tests for equality of proportions with continuity correction (2sEP) as implemented in the function ‘*prop.test*’ (library *stats*, (R Core Team 2016)).

Second, we studied the effects of PE copy number as well as the effects of its presence and absence. For each of two types of interspecific crosses for which we observed evidence of HD (i.e., ♀*sech* × ♂*sim* and ♀*mau* × ♂*sim*), we fitted a linear mixed model where the response of the model was whether a female showed evidence of dysgenesis or not, the type of cross (either conspecific or heterospecific) was a fixed effect, and the number of PE copies in the *D. simulans* genome was a continuous trait. Each of these datasets included interspecific data and intraspecific data (within *D. simulans*) from ♀ M × ♂ P crosses. We fitted a binomial regression using the function *glmer* in the library *lme4* (Bates *et al*., 2013). We included the interaction between these two effects to account for differences in HD caused by the number of PE copies and assessed its significance using a likelihood ratio test (function ‘*lrtest*’, R library *lmtest*; (Hothorn *et al*., 2011)) by comparing a model with the interaction and a model without it. The fully factorial model followed the form:

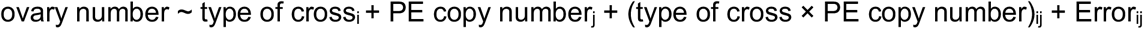

The model without the interaction followed the form:

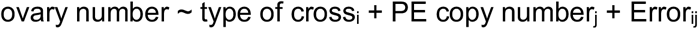

The goal of this comparison was to assess whether the slope of the regression differed between inter- and intra-specific crosses. To quantify the significance of each effect, we used a type III ANOVA in R (function ‘*Anova*’, library *car* (Fox and Sanford 2011; Fox and Weisberg 2019)). Since we used two metrics to estimate the number of PEs in each line genome (see above), we fitted sixteen linear models, one for each combination of cross, temperature, and PE copy number estimation (four types of crosses × two temperatures × two PE metrics).

### Male hybrid sterility

F_1_ hybrids between all interspecific crosses in the *simulans* species subgroup are sterile at 25°C (Turissini *et al*., 2018; Lachaise *et al*., 2018). We evaluated whether PE infection status affected this barrier. We made hybrid males as described above at two different temperatures: 25°C and 29°C. Four- to eight-day-old males from conspecific and heterospecific crosses were then lightly anesthetized with CO_2_. Their testes were then extracted with Dumont #5 forceps and mounted in chilled *Drosophila* Ringer’s solution (Cold Spring Harbor Protocols, Serrato-Capuchina *et al*., 2020). An average of five to ten pairs of testes were mounted per slide. Using a microscope, we scored whether each pair of testes had motile sperm within five minutes of having mounted the samples. For each interspecific genotype combination, we scored sperm motility for 100 males per cross (398 combinations: 39,800 total males).

## RESULTS

### TE survey in the *simulans* species complex

First, we explored whether the genomes of the three species from the *simulans* complex differed in their TE content. Figure 1A shows an LDA of the TEs that contribute to differences among genotypes (i.e., species and lines). We show the results when we allowed for up to 25% of a sequence to be missing in a TE to count it as present in that line. The first discriminant function, LD1, separates the three species (85% of the trace) more effectively than LD2 (15% of the trace). Figure 1B shows the loadings in LD1 for 27 TEs. PE and Het-A have the largest contributions to the discriminant function, but each of them contributes ~10% of the variance. Figure S1 shows the loadings for LD2, but this function is at least time five times less effective at separating species than LD1. These results suggest that the differences in TE content in the *simulans* species group are multifaceted and cannot be explained by a single type of TE. Figures S2-S3 show LDAs for a different, and more stringent, cutoff (necessitating 0% of the consensus sequence missing to count it as present in that line) to estimate TE copy number; regardless of the cutoff, *P*-element (PE) content differs between species in the *simulans* complex.

**FIGURE 1.**
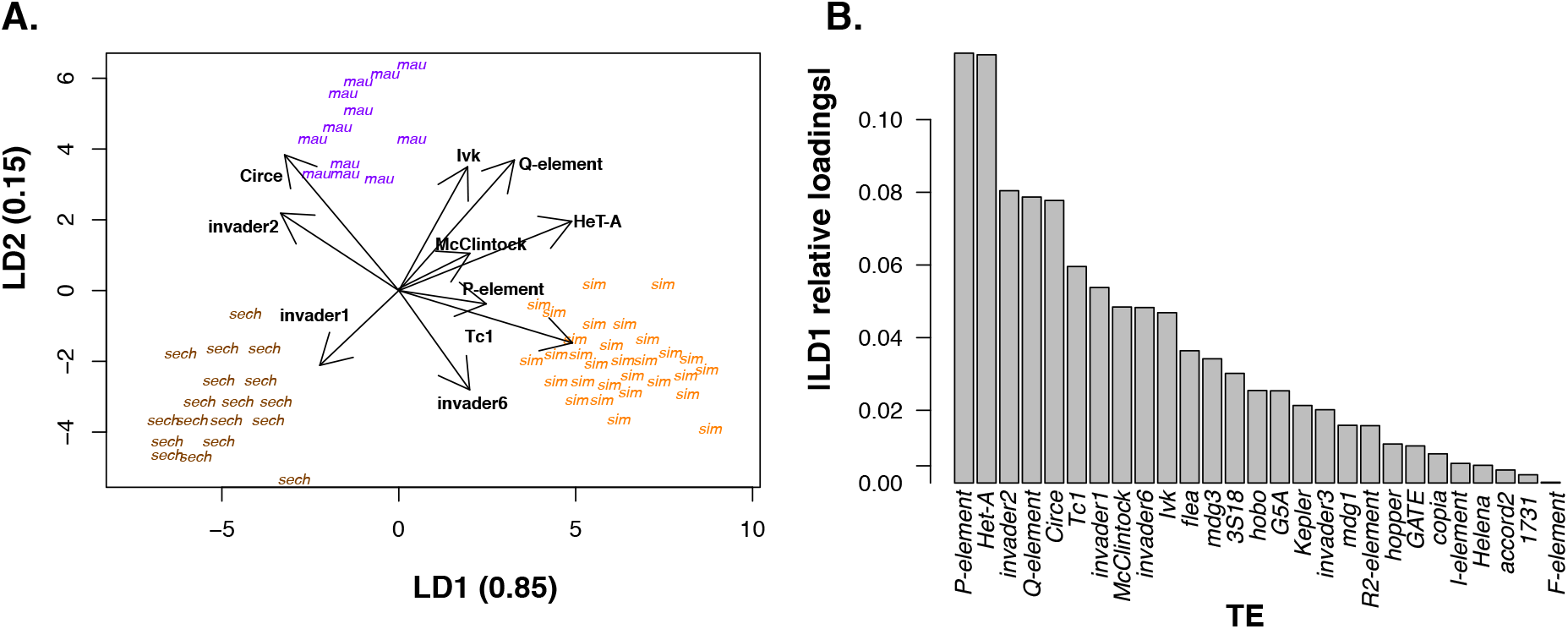
Linear discriminant analysis (LDA) of TEs present in the three species of the *simulans* species complex. **A.** Discriminant function scores of individuals from the three species of the *simulans* species complex. *D. sechellia*: brown, *D. mauritiana*: purple, *D. simulans*: orange. The axes show the proportion of trace (i.e., separation achieved) by each of the two discriminant functions. Arrows show the 10 TEs with the largest loadings **B.** Absolute value of the LD1 relative loadings by each of the 27 included TEs. PE and Het-A had the largest contributions to LD1. Figure S1 shows the LD2 relative loadings.

Next, we studied whether any TE was present in one of the species of the *simulans* clade but not in the other ones. Figure 2 shows the estimated number of copies for the twelve TEs that explain most of the variance in LD1 (listed in Figure 1B). Figures S4-S5 show the other 38 TEs. The majority of TEs were either present in all the three species—but in different copy numbers (Figure 2)—or absent in the three species (Figures S4-S5). This similarity reflects the close genealogical relationship between the triad of species. Different cutoffs had no effect on our results. Figure S6 shows boxplots for a more stringent cutoff to estimate TE copy number. Unlike *D. simulans*, which carry complete and incomplete copies of PEs (Kofler *et al*., 2015; Hill *et al*. 2016; Serrato-Capuchina *et al*., 2020), the latter two species have no traces of the PE sequence (i.e., no read coverage). The use of different cutoffs of sequence completeness to infer TE copy number had no effect on the inference of PE copy number (Figure 2A vs. Figure S6A). We verified this result using single-molecule sequencing and verified that neither *D. mauritiana* nor *D. sechellia* have complete—or incomplete—copies of PE.

**FIGURE 2.**
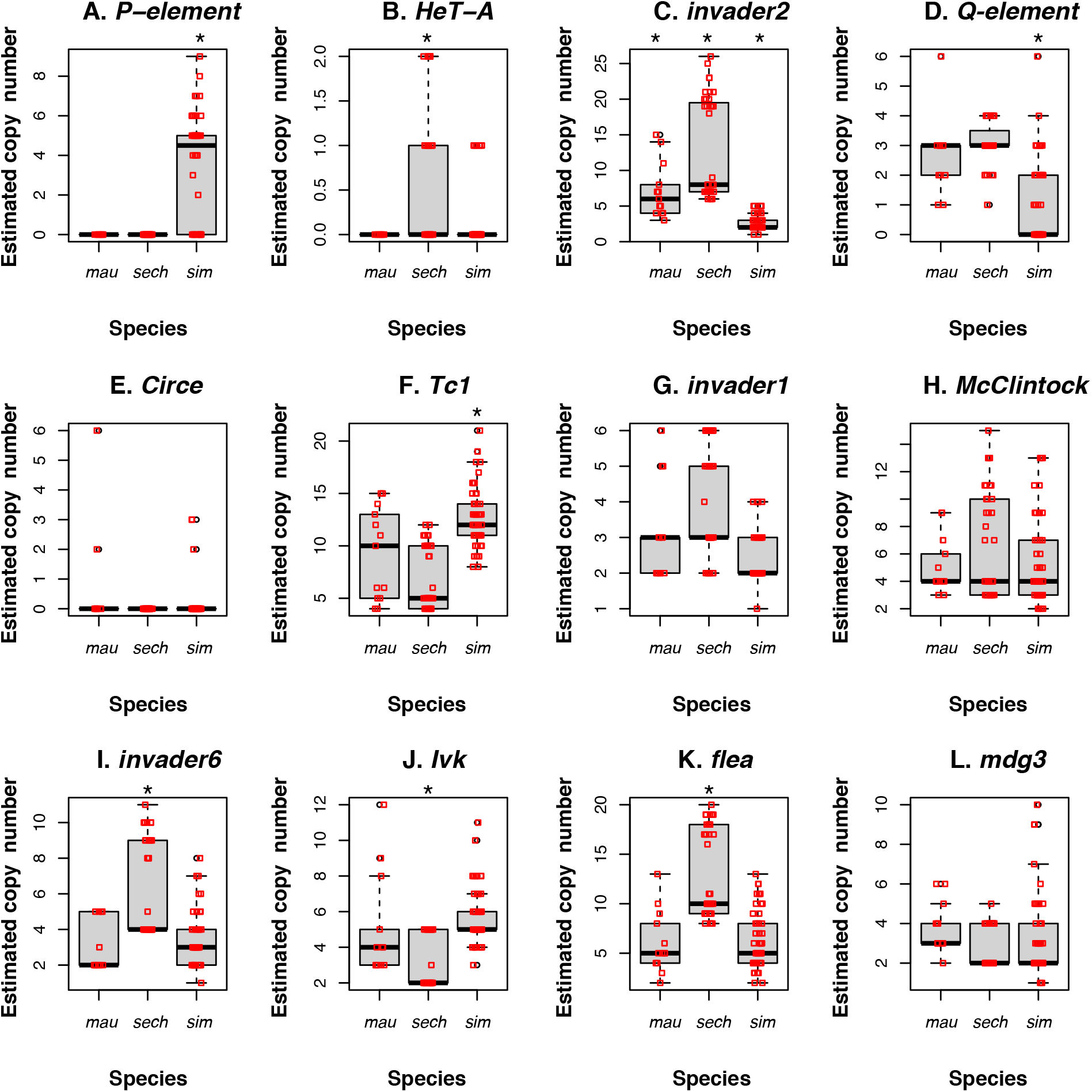
The abundance of TEs varies in the three species of the *simulans* complex. We filtered out copies in which at least 25% of the sequence of the TE was missing. We show the 12 TEs with the largest loadings in LD1; other TEs are shown in Figures S4-S5. Other filtering schemes are shown in Figures S6-S7. Each panel shows copy number for each of twelve TEs. Stars show instances where one species differed from the other two at a given TE copy-number (Tukey HSD tests). Please note that the *y*-axes differ across panels. The only TE present in one species but completely absent in the others is *P*-element.

Other TEs known to have an effect on HD (i.e. *I*-element and *hobo*) do have inferred copy numbers influenced by the choice of cutoff. When we used a 25% missing sequence cutoff, *I*-element and *hobo* were inferred to be present in the three species of the *simulans* complex. When we used a more stringent cutoff (0%), *I*-element was inferred to be absent in *D. sechellia* and *D. mauritiana; hobo* was inferred to be absent in *D. sechellia*. To resolve this discordance, we relied on long reads, which indicated that both *I*-element and *hobo* are indeed present in all the three species.

The species difference in the presence of PEs and *D. simulans*’ polymorphism for PE copy number and presence-absence make the *simulans* species complex ideal to study the effects of different types of transposable elements in RI in well-formed species. It is worth noting that two other TEs have been implicated in hybrid dysgenesis in *Drosophila: hobo* (Blackman *et al*. 1987; Ladevèze *et al*., 1998) and *I*-element (Picard *et al*., 1978; Orsi *et al*., 2010). Nonetheless, all lines from the *simulans* species complex carry *hobo* (Figure S4B) and *I*-elements (Figure S5R), so it is unlikely these two TEs, unlike PE, cause dysgenesis in interspecific hybrids.

### Effects of PE on female fecundity

PEs have been associated with reduced fecundity and higher rates of female sterility in within-species crosses in F1s (Bingham *et al*., 1982; Kidwell 1983b; Hill *et al*., 2016). PEs also differ in presence-absence between *D. simulans* and the other two species of the *simulans* complex (see above). We studied whether PEs had an effect on female hybrid fecundity. Specifically, we studied whether F1 hybrid females from ♀*mau* × ♂*sim* and ♀*sech* × ♂*sim* crosses were more likely to show atrophied ovaries, a proxy of hybrid dysgenesis if the fathers carried PEs. Figure 3 shows examples of progeny with atrophied ovaries from conspecific and heterospecific crosses. First, females from ♀M × ♂M crosses showed few females with atrophied gonads in both conspecific and heterospecific crosses (~0.1%, Table 1). Second, ♀M/♂P females from heterospecific crosses showed a higher rate of gonad atrophy than females from conspecific crosses at both temperatures (Figure 4, Table 1). ♀M/♂P F1 females from interspecific crosses were more likely to have atrophied ovaries than ♀M/♂P females from crosses within *D. simulans* (Table 1). Third, we found dysgenic females in all ♀M ×♂ P crosses at both temperatures but, as expected (Kidwell and Novy 1979), the frequency of gonad atrophy in ♀M × ♂ P crosses is higher at 29°C than at 25°C in conspecific crosses (2sEP: X^2^ = 159.37, df = 1, P < 1 × 10^−10^) and heterospecific crosses (2sEP _*mau* × *sim*_: χ^2^= 122.76, df = 1, P < 1 × 10^−10^; 2sEP_*sech* × *sim*_: χ^2^= 18.593, df = 1, P = 1.618 × 10^−5^; Figure 4A vs. 4C and Figure 4B vs. 4D). These results suggest that PE-induced hybrid dysgenesis—at least in the form of ovary atrophy—is more pronounced in heterospecific than in conspecific crosses.

**FIGURE 3.**
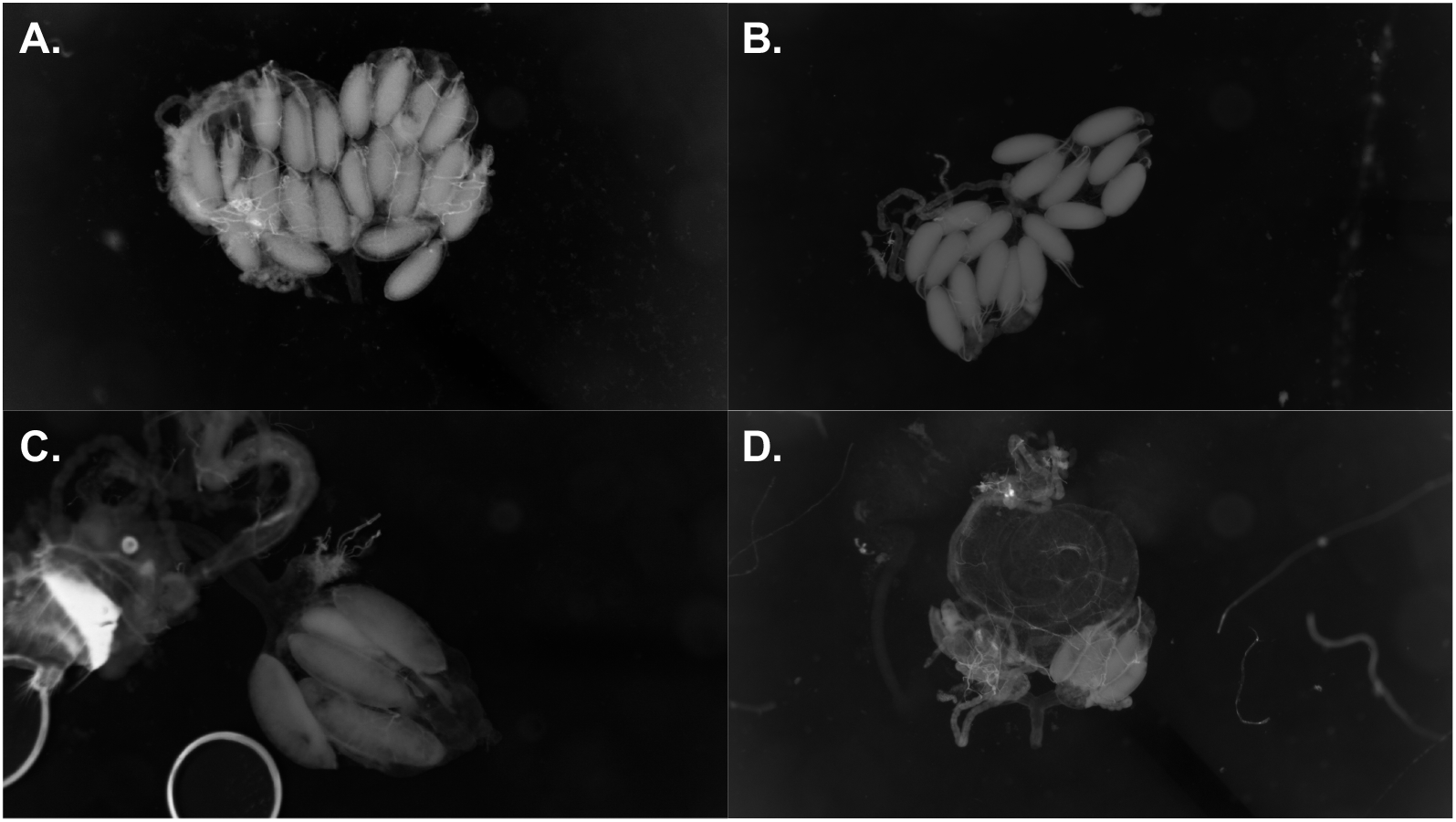
Examples of non-dysgenic and dysgenic female reproductive tracts from conspecific and heterospecific F1s. **A.** Non-dysgenic ♀ M/♂M *D. simulans/D. simulans* F1 female. **B.** Non-dysgenic ♀ M/♂M F1 *D. mauritiana* female. **C.** Dysgenic ♀M/♂P *D. simulans/D. simulans* F1 female. **D.** Dysgenic ♀M/♂P *D. mauritiana/D. simulans* F1 female. Both ♀M/♂P females (**C** and **D**) have only one functional ovary.

**TABLE 1.** ♀M/♂P females from heterospecific crosses are more likely to be dysgenic than ♀M/♂P females from conspecific crosses. Females with atrophied ovaries are more common in heterospecific than in conspecific crosses in this direction of the cross. Each row shows a type of cross. Temp (°C): Temperature (°C); CI: Binomial confidence interval. We used proportion tests for all comparisons and adjusted P-values with a Bonferroni correction.

|  |  | ♀M/♂P |  |  | ♀M/♂M |  |  | Proportion test |  |  |
| --- | --- | --- | --- | --- | --- | --- | --- | --- | --- | --- |
|  | Temp (°C) | Total | Mean | CI | Total | Mean | CI | X<sup>2</sup> | df | P |
| ♀*sim* ×<br>♂*sim* | 25 | 2,100 | 0.982 | [0.975, 0.987] | 1,960 | 0.999 | [0.996, 1.000] | 28.574 | 1 | <0.0001 |
| ♀*mau* ×<br>♂*sim* | 25 | 980 | 0.851 | [0.827, 0.872] | 980 | 1.000 | [0.996, 1.000] | 44.175 | 1 | <0.0001 |
| ♀*sech* ×<br>♂*sim* | 25 | 980 | 0.821 | [0.796, 0.844] | 980 | 0.986 | [0.976, 0.992] | 18.593 | 1 | <0.0001 |
| ♀*sim* ×<br>♂*sim* | 29 | 2,100 | 0.829 | [0.812, 0.844] | 1,960 | 0.988 | [0.982, 0.992] | 299.22 | 1 | <0.0001 |
| ♀*mau* ×<br>♂*sim* | 29 | 980 | 0.728 | [0.699, 0.754] | 980 | 1.000 | [0.996, 1.000] | 44.175 | 1 | <0.0001 |
| ♀*sech* ×<br>♂*sim* | 29 | 980 | 0.740 | [0.711, 0.766] | 980 | 1.000 | [0.996, 1.000] | 18.593 | 1 | <0.0001 |

**FIGURE 4.**
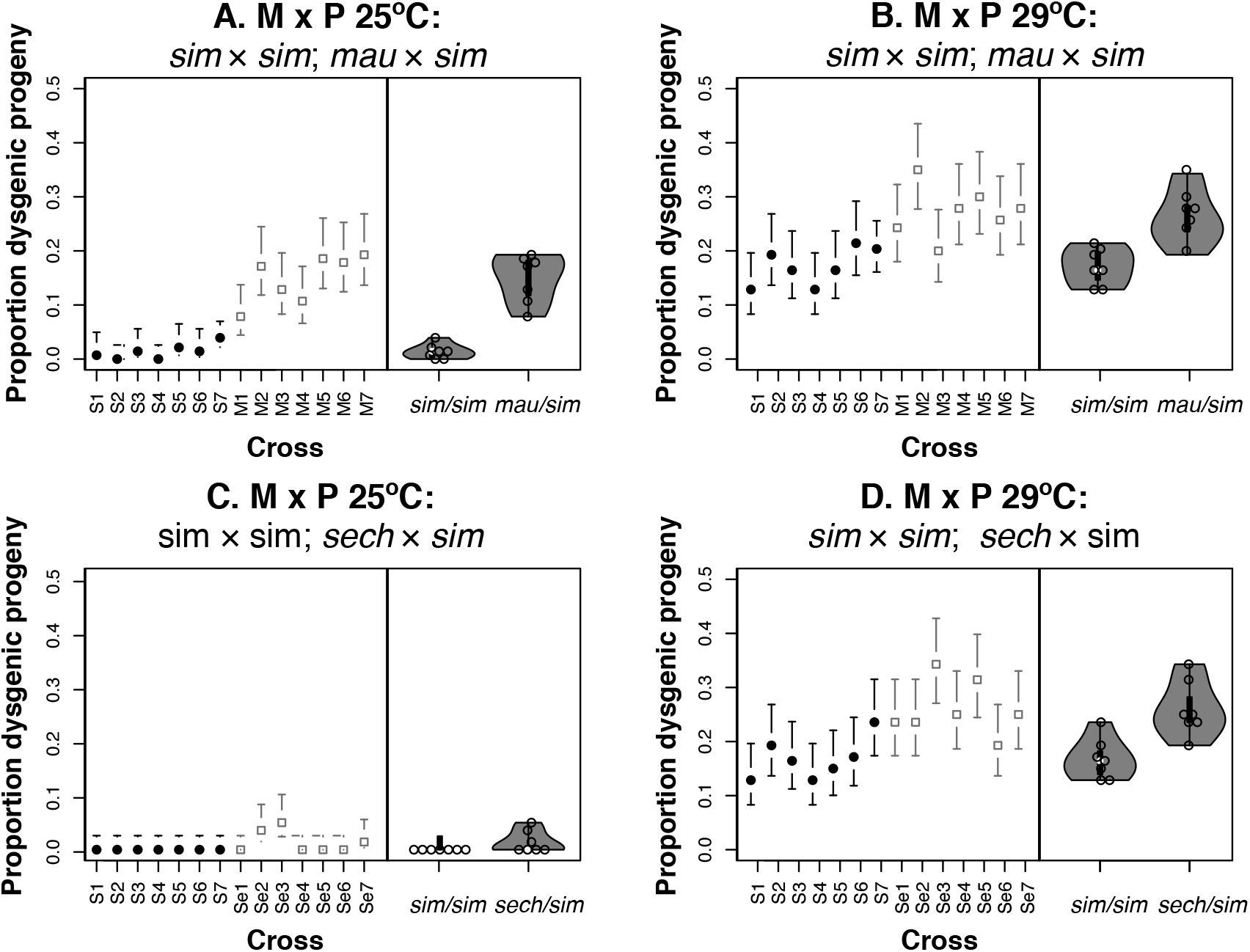
The proportion of dysgenic progeny is higher in heterospecific than in conspecific ♀M/♂P crosses. Left panels: Solid black circles represent conspecific crosses within *D. simulans* (♀ *D. simulans* × ♂ *D. simulans*); open squares represent interspecific crosses (either ♀ *D. mauritiana* × ♂ *D. simulans* or (♀ *D. sechellia* × ♂ *D. simulans*). Bars show the Bayesian confidence interval of the proportion. Right panels: Violin plots showing the distribution of the means for each maternal line (shown in left panels) for each type of cross (conspecific vs. heterospecific). **A.** ♀ *sim* M × ♂ *sim* P and ♀ *mau* M × ♂ *sim* P at 25°C. **B.** ♀ *sim* M × ♂ *sim* P and ♀ *mau* M × ♂ *sim* P at 29°C. **C.** ♀ sim M × ♂ *sim* P and ♀ *sech* M × ♂ *sim* P at 25°C. **D.** ♀ *sim* M × ♂ *sim* P and ♀ *sech* M × ♂ *sim* P at 29°C. S1: Md06, S2: Md105, S3: Md106, S4: Md199, S5: NS05, S6: NS113, S7: NS40; *D. mauritiana* lines: M1: R23, M2: R31, M3: R32, M4: R42, M5: R50, M6: R61, M7: R8; *D. sechellia* lines: Se1: Anro71, Se2: Denis124, Se3: Denis72, Se4: DenisNF100, Se5: DenisNF13, Se6: LD11, Se7: LD16.

An additional prediction of PE-induced HD is that F1 females from ♀P × ♂M crosses (♀P/♂M) should show similar ovary counts to F1 ♀M/♂M females in conspecific and heterospecific crosses. Regardless of the cross and the temperature, the vast majority of conspecific and hybrid females with a *D. simulans* mother had two ovaries (Table S4). As predicted, ♀P/♂M F1s from either conspecific or heterospecific crosses are non-dysgenic.

### PE copy number effect on female fecundity

PE-copy number in the paternal genome is correlated with the strength of hybrid dysgenesis; paternal lines with more PEs cause stronger hybrid dysgenesis in conspecific crosses ((Bingham *et al*., 1982; Hill *et al*., 2016; Srivastav and Kelleher 2017; Serrato-Capuchina *et al*., 2020) but see (Bergman *et al*., 2017; Srivastav *et al*., 2019)). We compared the effect of PE copy number on the number of ovaries produced in conspecific and heterospecific females from M × P crosses (♀M/♂P). The results at the two temperatures, 25°C and 29°C, were qualitatively similar (Figure 2; Table 2). ♀M/♂P (*D. mauritiana × D. simulans*) F1 females are less likely to be fecund than ♀M/♂P (*D. simulans × D. simulans*) F1 females at both 25°C and 29°C. As the PE copy number in the paternal genome increases, the likelihood of an ♀M/♂P F1 female being fecund decreases (odds ratios in Table 2). The rate of decrease in female fecundity as PE increases was similar in heterospecific and conspecific crosses (interaction in Table 2). We found a similar result when we compared ♀M/♂P (*D. sechellia × D. simulans*) and ♀M/♂P (*D. simulans × D. simulans*) females. F1 ♀M/♂P females from heterospecific crosses less likely to be fecund than F1 females from conspecific crosses to be dysgenic at both 25°C and 29°C (odds ratios in Table 2). The PE copy number effect was also significant, indicating the existence of a negative relationship between paternally inherited PEs and ovary number in the female ♀M/♂P progeny (Table 2). The interaction term (PE copy × cross) in the comparison at 29°C was significant (Likelihood Ratio test; χ^2^ = 7.605, df = 1, P = 5.821 × 10^−3^). This result suggests that the slope of the decrease in fecundity as PE copy number increases was slightly more pronounced in ♀M/♂P heterospecific crosses than in ♀M/♂P conspecific crosses (Slope _*sech* × *sim*_ — slope _*sim* × *sim*_ = −0.083, SD = 0.031, Z = −2.721, P= 6.51 × 10^−3^). Results for all crosses were identical when we used a previous inference of PE copy number ((Serrato-Capuchina *et al*. 2020), Table S5).

**TABLE 2.** PE copy number in the paternal genome affects the number of functional ovaries in conspecific and heterospecific produced F1 females. Each row shows the results for each cross at one of two temperatures (°C). The linear models for each combination of cross and temperature were binomial models fitted to compare the effect of paternally inherited PEs on conspecific and hybrid F1 females. The effect ‘cross’ had two levels (*D. simulans × D. simulans* and a heterospecific cross) while the PE copies effect was continuous. For the cross effect, we show the log-odds ratio of being non-dysgenic between F1 (♀*sim*/♂*sim*) and *F1*(♀*mau*/♂*sim*) (or *F1*(♀*sech*/♂*sim*)).

| Cross | °C | PE Copies |  |  | Cross |  |  | PE copies × cross |  |
| --- | --- | --- | --- | --- | --- | --- | --- | --- | --- |
|  |  | Log odds ratio CI | F | P | Odds ratio CI | F | P | LRT, X<sup>2</sup><sub>1</sub> | P |
| ♀*sech* × ♂*sim* | 25 | [-0.286, -0.210] | 161.830 | < 1×10<sup>-5</sup> | [2.049, 3.021] | 103.340 | < 1×10<sup>-5</sup> | 0.404 | 0.525 |
| ♀*sech* × ♂*sim* | 29 | [-0.313, -0.256] | 198.724 | < 1×10<sup>-5</sup> | [0.437, 0.952] | 25.827 | < 1×10<sup>-5</sup> | 7.605 | 5.821×10<sup>-3</sup> |
| ♀*mau* × ♂*sim* | 25 | [-0.278, -0.195] | 125.879 | < 1×10<sup>-5</sup> | [1.710, 2.941] | 59.021 | < 1×10<sup>-5</sup> | 2.405 | 0.121 |
| ♀*mau* × ♂*sim* | 29 | [-0.344, -0.286] | 438.600 | < 1×10<sup>-5</sup> | [0.327, 0.865] | 21.009 | < 1×10<sup>-5</sup> | 0.102 | 0.7498 |

### Effects of PE on male fecundity

We scored sperm motility in hybrid males involving *D. sechellia*, or *D. mauritiana*, mothers, and P *D. simulans* fathers. All F1 males, from all interspecific crosses (N=100 per cross), were completely sterile and produced no sperm, regardless of the identity of the cross. The PE status of *D. simulans* did not affect hybrid male fertility.

## DISCUSSION

*P*-elements (PEs) in *Drosophila* cause hybrid dysgenesis and remain one of the best-understood cases of how TEs interact with their host genome (reviewed in (Kelleher 2016)). In this report, we studied the role of PEs in RI between three closely related species of the *simulans* species complex by crossing *D. simulans* lines that varied in whether they carry PEs to two sister species that do not carry PEs. We find that PE status of the father (and the number of PE copies) affects F1 female fertility in interspecific crosses. HD in F1 ♀M/♂P females is stronger in hybrid females than in within-species females, suggesting that hybrids are more susceptible to the effects of HD than pure-species individuals.

The importance of TEs in the speciation process remains largely unexplored (Watson and Demuth 2013; Serrato-Capuchina and Matute 2018). In the case reported here, PEs cause HD in both pure species and hybrid F1 females. F1 ♀M/♂P females should suffer the deleterious effects of PEs because they do not have the maternally deposited piRNAs that confer protection against TE misregulation (Michalak 2009; Majumdar and Rio 2015; Kelleher 2016). This should be true of ♀M/♂P F1s from either pure-species or interspecific crosses. Our results show that even though PEs induce female HD in conspecific and interspecific ♀M/♂P F1s, the magnitude of HD is moderately stronger in interspecific hybrids than in within-species ♀M/♂P F1s. There must, therefore, be genetic interactions between the two parental genomes and PEs that affect the magnitude of HD in F1s. There is precedent supporting this hypothesis. In interspecific crosses involving a *D. lummei* mother and a *D. virilis* father, interspecific hybrids show exacerbated gonadal dystrophy (a proxy of HD) when the father has the TE, *Penelope*. Genetic analyses suggest that *Penelope* interacts with other autosomal and sex-linked elements by modifying the penetrance of hybrid incompatibilities (Castillo and Moyle, 2019).

TE expression is regularly higher in hybrids compared than in pure species. In hybrid rice (Shan *et al*., 2005) and hybrid sunflowers (Ungerer *et al*., 2006) there is extensive TE mobilization and proliferation, indicating that these defects might be common among seed plant hybrids. In *Drosophila*, TEs are often overexpressed in *Drosophila* hybrids (Kelleher *et al*., 2012; Lopez-Maestre *et al*., 2017; Romero-Soriano *et al*., 2017). Misregulation of TEs in hybrids might be related to the divergence of PIWI and piRNA pathways across species, either by natural selection, genetic conflict, or systems drift (i.e., divergence in the molecular underpinnings of homologous traits not due directly to selection, reviewed in (True and Haag 2001)). Genes from the PIWI pathway show signatures of positive selection since the divergence between *D. simulans* and *D. melanogaster* ((Vermaak *et al*., 2005; Heger and Ponting 2007; Obbard *et al*., 2009; Kolaczkowski *et al*., 2011; Lee and Langley 2012; Simkin *et al*., 2013; Wang *et al*., 2020) reviewed in (Blumenstiel *et al*., 2016)). At least three piRNA-interacting genes, *aubergine* (Kelleher *et al*., 2012; Wang *et al*., 2020)*, Armitage* and *Spindle-E* (Wang *et al*., 2020) have functionally diverged between *D. simulans* and *D. melanogaster*. Functional divergence in these proteins leads to off-target effects on host transcripts in *D. melanogaster* transgenics carrying the *D. simulans* allele. This divergence might also play a role in reduced hybrid fitness. A decrease in genomic autoimmunity in hybrids caused by adaptive divergence of genes in the PIWI pathway might join PE misexpression as a cause of PE-induced HD in interspecific hybrids and conspecific F1s.

The relatively recent invasions of PEs in both *D. simulans* and *D. melanogaster* seem to be explained by horizontal gene transfer through an unknown vehicle, which poses the question of how often TEs move between species and establish successfully in the host genome. A similarly intriguing question is why PEs have not been found in *D. mauritiana* and *D. sechellia*. In both of these island endemics, there is evidence for interspecific introgression with *D. simulans* (Brand *et al*., 2013; Garrigan *et al*., 2012; Meiklejohn *et al*., 2018), and even though alleles have crossed those species boundaries, PEs have not been transferred from *D. simulans* to *D. mauritiana*. Hybridization can be an effective mechanism of TE transfer across species boundaries (Vela *et al*., 2014; Heil *et al*., 2020). One possibility is that introgression is ancient and precedes the invasion of PEs in *D. simulans*. The case of *D. sechellia* is even more puzzling. Not only do *D. sechellia* and *D. simulans* show signatures of reciprocal introgression (Schrider *et al*., 2018), these two species form a contemporary hybrid zone in the central islands of the Seychelles archipelago (Matute and Ayroles 2014). This indicates that, at present, there is ample opportunity for gene exchange—and for interspecific transfer of PEs. Alternatively, the *D. sechellia* and *D. mauritiana* genomes could be more refractory to the invasion of PEs than *D. simulans*. The evolutionary dynamics of TE copy number in admixed lineages can differ strongly even among evolutionarily closely related hybrids (Henault *et al*., 2020). Experimental infections with PEs showed the potential of *D. simulans* to host PEs (Daniels *et al*., 1989). The same type of experiments could reveal if PEs are less successful at invading some genomes than others, and this, in turn, could explain the absence of PEs in the *D. sechellia* and *D. mauritiana* genomes.

The relative importance of different mechanisms of successful genome invasion by TEs (horizontal gene transfer vs. introgression) remains largely unexplored. It is likely that multiple mechanisms play a role in the transmission of PEs, and only a robust systematic assessment of the relative frequency of PEs in different species of *Drosophila* and the putative vectors will reveal the relative importance of each mechanism in the spread of PEs within and between species. Our results show that the dynamics of PEs, and possibly of TEs in general, should not only be addressed with a lens on the fitness effects that PEs have on within-species crosses, but also on the effects that they might have on interspecific hybrids.

## REFERENCES

Barbash, D. A., and M. Ashburner. 2003. A novel system of fertility rescue in *Drosophila* hybrids reveals a link between hybrid lethality and female sterility. Genetics 163: 217–226.

Bates, D., M. Maechler, B. Bolker, and S. Walker. 2013. lme4: Linear mixed-effects models using Eigen and S4. R Packag. version. 1.1-7

Bergman, C. M., S. Han, M. G. Nelson, V. Bondarenko, and I. Kozeretska. 2017. Genomic analysis of P elements in natural populations of *Drosophila melanogaster*. PeerJ, 5: p.e3824. doi: 10.7717/peerj.3824.

Bingham, P. M., M. G. Kidwell, and G. M. Rubin. 1982. The molecular basis of P-M hybrid dysgenesis: The role of the *P* element, a *P*-strain-specific transposon family. Cell, 29: 995–1004. doi: 10.1016/0092-8674(82)90463-9.

Blackman, R. K., R. Grimaila, M. Macy, D. Koehler, and W. M. Gelbart. 1987. Mobilization of *hobo* elements residing within the decapentaplegic gene complex: Suggestion of a new hybrid dysgenesis system in *Drosophila melanogaster*. Cell, 49: 497–505.doi: 10.1016/0092-8674(87)90452-1.

Blumenstiel, J. P., A. A. Erwin, and L. W. Hemmer. 2016. What drives positive selection in the Drosophila piRNA machinery? The genomic autoimmunity hypothesis. The Yale journal of biology and medicine, 89: 499.

Brand, C. L., S. B. Kingan, L. Wu, and D. Garrigan. 2013. A selective sweep across species boundaries in *Drosophila*. Mol. Biol. Evol., 30: 2177–2186. doi: 10.1093/molbev/mst123.

Brideau, N. J., H. A. Flores, J. Wang, S. Maheshwari, X. Wang, and D. A. Barbash. 2006. Two Dobzhansky-Muller Genes interact to cause hybrid lethality in *Drosophila*. Science 314:1292–1295.

Brookfield, J. F., and R. M. Badge. 1997. Population genetics models of transposable elements. Genetica 100:281–294.

Brookfield, J. F. Y., E. Montgomery, and C. H. Langley. 1984. Apparent absence of transposable elements related to the *P* elements of *D. melanogaster* in other species of *Drosophila*. Nature, 310: 330–332. doi: 10.1038/310330a0.

Castillo, D. M., and L. C. Moyle. 2012. Evolutionary implications of mechanistic models of TE-mediated hybrid incompatibility. Int. J. Evol. Biol., doi: 10.1155/2012/698198.

Clark, J. B., T. K. Altheide, M. J. Schlosser, and M. G. Kidwell. 1995. Molecular evolution of *P* transposable elements in the genus *Drosophila*. I. The *saltans* and *willistoni* species groups. Mol. Biol. Evol. 12:902–13.

Comeault, A. A., A. Serrato Capuchina, D. A. Turissini, P. J. McLaughlin, J. R. David, and D. R. Matute. 2017. A nonrandom subset of olfactory genes is associated with host preference in the fruit fly *Drosophila orena*. Evol. Lett. 1: 73–85.

Daniels, S. B., A. Chovnick, and M. G. Kidwell. 1989. Hybrid dysgenesis in *Drosophila simulans* lines transformed with autonomous *P* elements. Genetics. 121: 281–291.

Daniels, S. B., K. R. Peterson, L. D. Strausbaugh, M. G. Kidwell, and A. Chovnik. 1990. Evidence for horizontal transmission of the *P* transposable element between *Drosophila* species. Genetics. 124: 339–355.

Davis, A. W., J. Roote, T. Morley, and K. Sawamura. 1996. Rescue of hybrid sterility in crosses between *D. melanogaster* and *D. simulans*. Nature, 380: 157–159.

De Setta, N., A. P. P. Costa, F. R. Lopes, M. A. Van Sluys, and C. M. A. Carareto. 2007. Transposon display supports transpositional activity of *P* elements in species of the *saltans* group of *Drosophila*. J. Genet., 86: 37. doi: 10.1007/s12041-007-0005-z.

Dion-Côté, A.-M., S. Renaut, E. Normandeau, and L. Bernatchez. 2014. RNA-seq reveals transcriptomic shock involving transposable elements reactivation in hybrids of young lake whitefish species. Mol. Biol. Evol. 31:1188–1199.

Dion-Côté, A. M., and D. A. Barbash. 2017. Beyond speciation genes: an overview of genome stability in evolution and speciation. Current opinion in genetics & development, 47: 17–23.

Dorai-Raj, S. 2015. Package “binom”. Binomial Confidence Intervals For Several Parameterizations. CRAN R Proj.

Engels, W. R., and C. R. Preston. 1980. Components of hybrid dysgenesis in a wild population of *Drosophila melanogaster*. Genetics 95:111–128.

Fox, J., and W. Sanford. 2011. car: An R Companion to Applied Regression (R package).

Fox, J., and S. Weisberg. 2019. An {R} Companion to Applied Regression, (top).

Garrigan, D., S. B. Kingan, A. J. Geneva, P. Andolfatto, A. G. Clark, K. R. Thornton, and D. C. Presgraves. 2012. Genome sequencing reveals complex speciation in the *Drosophila simulans* clade. Genome Res. 22:1499–1511.

Ginzburg, L. R., P. M. Bingham, and S. Yoo. 1984. On the theory of speciation induced by transposable elements. Genetics 107:331–341.

Heger, A., and C. P. Ponting. 2007. Evolutionary rate analyses of orthologs and paralogs from 12 *Drosophila* genomes. Genome Res., 17: 1837–1849. doi: 10.1101/gr.6249707.

Heil, C. S., K. Patterson, A. S.-M. Hickey, E. Alcantara, and M. J. Dunham. 2020. Transposable element mobilization in interspecific yeast hybrids. bioRxiv, doi: 10.1101/2020.06.16.155218.

Hill, T., C. Schlotterer, and A. J. Betancourt. 2016. Hybrid dysgenesis in *Drosophila simulans* associated with a rapid invasion of the *P*-element. PLOS Genetics 12: e1005920.

Hothorn, A. T., A. Zeileis, G. Millo, and D. Mitchell. 2011. Package ‘lmtest.’ R News.

Hothorn, T., F. Bretz, and P. Westfall. 2008. Simultaneous Inference in Genneral Parametric Models. Biometrical J. 50: 346–363.

Ishikawa, R., T. Ohnishi, Y. Kinoshita, M. Eiguchi, N. Kurata, and T. Kinoshita. 2011. Rice interspecies hybrids show precocious or delayed developmental transitions in the endosperm without change to the rate of syncytial nuclear division. Plant J. 65:798–806.

Kelleher, E. S. 2016. Reexamining the P-element invasion of *Drosophila melanogaster* through the lens of piRNA silencing. Genetics, 203: 1513–1531.

Kelleher, E. S., N. B. Edelman, and D. A. Barbash. 2012. *Drosophila* interspecific hybrids phenocopy piRNA-pathway mutants. PLoS Biol. 10: e1001428

Kidwell, M. G. 1983a. Evolution of hybrid dysgenesis determinants in *Drosophila melanogaster*. Proc. Natl. Acad. Sci. U. S. A. 80: 1655–9.

Kidwell, M. G. 1983b. Hybrid dysgenesis in *Drosophila melanogaster*: Factors affecting chromosomal contamination in the P-M system. Genetics, 104: 317–341.

Kidwell, M. G. 1985. Hybrid dysgenesis in *Drosophila melanogaster*: Nature and inheritance of P element regulation. Genetics. 111: 337–350.

Kidwell, M. G., and J. F. Kidwell. 1975. Cytoplasm-chromosome interactions in *Drosophila melanogaster*. Nature, 253: 755–756. doi: 10.1038/253755a0.

Kidwell, M. G., J. F. Kidwell, and J. A. Sved. 1977. Hybrid dysgenesis in *Drosophila melanogaster*: a syndrome of aberrant traits including mutation, sterility and male recombination. Genetics 86: 813–833.

Kidwell, M. G., and J. B. Novy. 1979. Hybrid dysgenesis in *Drosophila melanogaster*: Sterility resulting from gonadal dysgenesis in the P-M system. Genetics, 92: 1127–1140.

Kofler, R., T. Hill, V. Nolte, A. J. Betancourt, and C. Schlötterer. 2015. The recent invasion of natural *Drosophila simulans* populations by the *P*-element. Proc. Natl. Acad. Sci. 112: 6659–6663.

Kolaczkowski, B., D. N. Hupalo, and A. D. Kern. 2011. Recurrent adaptation in RNA interference genes across the *Drosophila* phylogeny. Mol. Biol. Evol., 28: 1033–1042. doi: 10.1093/molbev/msq284.

Lachaise, D., J. R. David, F. Lemeunier, L. Tsacas, and M. Ashburner. 1986. The reproductive relationships of *Drosophila sechellia* with *D. mauritiana, D. simulans*, and *D. melanogaster* from the Afrotropical region. Evolution. 40: 262–271.

Ladevèze, V., S. Aulard, N. Chaminade, G. Périquet, and F. Lemeunier. 1998. *Hobo* transposons causing chromosomal breakpoints. Proc. Biol. Sci. 265:1157–9.

Lee, Y. C. G., and C. H. Langley. 2012. Long-term and short-term evolutionary impacts of transposable elements on *Drosophila*. Genetics, 192: 1411–1432. doi: 10.1534/genetics.112.145714.

Li, H. 2016. Minimap and miniasm: Fast mapping and de novo assembly for noisy long sequences. Bioinformatics, 32: 2103–2110. doi: 10.1093/bioinformatics/btw152.

Li, H. 2018. Minimap2: Pairwise alignment for nucleotide sequences. Bioinformatics, 34: 3094–3100. doi: 10.1093/bioinformatics/bty191.

Lopez-Maestre, H., E. A. G. Carnelossi, V. Lacroix, N. Burlet, B. Mugat, S. Chambeyron, C. M. A. Carareto, and C. Vieira. 2017. Identification of misexpressed genetic elements in hybrids between *Drosophila*-related species. Sci. Rep., 7: 1–13. doi: 10.1038/srep40618.

Majumdar, S., and D. C. Rio. 2015. *P* Transposable elements in *Drosophila* and other Eukaryotic organisms. Chapter 33, pp. 727–752 in Mobile DNA III, Edited by: Chandler, M.; Gellert, M.; Lambowitz, A. M.; Rice, P. A.; Sandmeyer, S.B. Washington, DC: ASM Press

Matute, D. R., and J. F. Ayroles. 2014. Hybridization occurs between *Drosophila simulans* and *D. sechellia* in the Seychelles archipelago. J. Evol. Biol. 27:1057–1068.

Matute, D. R., A. A. Comeault, E. Earley, A. Serrato-Capuchina, D. Peede, A. Monroy-Eklund, W. Huang, C. D. Jones, T. F. C. Mackay, and J. A. Coyne. 2020. Rapid and predictable evolution of admixed populations between two *Drosophila* species pairs. Genetics 214:211–30. doi: 10.1534/genetics.119.302685.

Matute, D. R., and J. A. Coyne. 2010. Intrinsic reproductive isolation between two sister species of *Drosophila*. Evolution 64: 903–920.

McClintock, B. 1953. Induction of Instability at selected loci in maize. Genetics, 38: 579–599.

McQuillan, M. A., T. C. Roth, A. V. Huynh, and A. M. Rice. 2018. Hybrid chickadees are deficient in learning and memory. Evolution 72: 1155–1164. doi: 10.1111/evo.13470.

Meiklejohn, C. D., E. L. Landeen, K. E. Gordon, T. Rzatkiewicz, S. B. Kingan, A. J. Geneva, J. P. Vedanayagam, C. A. Muirhead, D. Garrigan, D. L. Stern, and D. C. Presgraves. 2018. Gene flow mediates the role of sex chromosome meiotic drive during complex speciation. Elife, 7, p.e35468.

Metcalfe, C. J., K. V. Bulazel, G. C. Ferreri, E. Schroeder-Reiter, G. Wanner, W. Rens, C. Obergfell, M. D. B. Eldridge, and R. J. O’Neill. 2007. Genomic instability within centromeres of interspecific marsupial hybrids. Genetics 177: 2507–2517.

Michalak, P. 2009. Epigenetic, transposon and small RNA determinants of hybrid dysfunctions. Heredity (Edinb). 102: 45–50.

Mizrokhi, L. J., and A. M. Mazo. 1990. Evidence for horizontal transmission of the mobile element *jockey* between distant *Drosophila* species. Proc. Natl. Acad. Sci. 87: 9216–9220.

Moyle, L. C., M. S. Olson, and P. Tiffin. 2004. Patterns of reproductive isolation in three angiosperm genera. Evolution 58: 1195–1208. doi: 10.1111/j.0014-3820.2004.tb01700.x.

Nolte, A. W., S. Renaut, and L. Bernatchez. 2009. Divergence in gene regulation at young life history stages of whitefish (*Coregonus* sp.) and the emergence of genomic isolation. BMC Evol. Biol. 9: 59.

Obbard, D. J., J. J. Welch, K. W. Kim, and F. M. Jiggins. 2009. Quantifying adaptive evolution in the *Drosophila* immune system. PLoS Genet., 5: e1000698. doi: 10.1371/journal.pgen.1000698.

Orr, H. A. 2005. The genetic basis of reproductive isolation: Insights from *Drosophila*. Proc. Natl. Acad. Sci. 102: 6522–6526.

Orr, H. A., and D. C. Presgraves. 2000. Speciation by postzygotic isolation: Forces, genes and molecules. BioEssays 22: 1085–1094.

Orsi, G. A., E. F. Joyce, P. Couble, K. S. McKim, and B. Loppin. 2010. *Drosophila* I-R hybrid dysgenesis is associated with catastrophic meiosis and abnormal zygote formation. J. Cell Sci., 123: 3515–3524. doi: 10.1242/jcs.073890.

Pélissier, T., S. Tutois, J. M. Deragon, S. Tourmente, S. Genestier, and G. Picard. 1995. *Athila*, a new retroelement from *Arabidopsis* thaliana. Plant Mol. Biol. 29: 441–452.

Pélissier, T., S. Tutois, S. Tourmente, J. M. Deragon, and G. Picard. 1996. DNA regions flanking the major *Arabidopsis thaliana* satellite are principally enriched in *Athila* retroelement sequences. Genetica 97: 141–51.

Picard, G., J. C. Bregliano, A. Bucheton, J. M. Lavige, A. Pelisson, and M. G. Kidwell. 1978. Non-mendelian female sterility and hybrid dysgenesis in *Drosophila melanogaster*. Genet. Res., 32: 275–287. doi: 10.1017/S0016672300018772.

R Core Team. 2016. R Development Core Team.

Ramsey, J., H. D. Bradshaw, and D. W. Schemske. 2003. Components of reproductive isolation between the monkeyflowers *Mimulus lewisii* and *M. cardinalis* (Phrymaceae). Evolution, 57: 1520–1534. doi: 10.1111/j.0014-3820.2003.tb00360.x.

Rogers, R. L., J. M. Cridland, L. Shao, T. T. Hu, P. Andolfatto, and K. R. Thornton. 2014. Landscape of standing variation for tandem duplications in *Drosophila yakuba* and *Drosophila simulans*. Mol. Biol. Evol. 31: 1750–1766.

Romero-Soriano, V., L. Modolo, H. Lopez-Maestre, B. Mugat, E. Pessia, S. Chambeyron, C. Vieira, and M. P. Garcia Guerreiro. 2017. Transposable element misregulation is linked to the divergence between parental piRNA pathways in *Drosophila* hybrids. Genome Biol. Evol., 9: 1450–1470. doi: 10.1093/gbe/evx091.

Rosenblum, E. B., B. A. J. Sarver, J. W. Brown, S. Des Roches, K. M. Hardwick, T. D. Hether, J. M. Eastman, M. W. Pennell, and L. J. Harmon. 2012. Goldilocks meets Santa Rosalia: an ephemeral speciation model explains patterns of diversification across time scales. Evol. Biol., 39: 255–261. doi: 10.1007/s11692-012-9171-x.

Rubin, G. M., M. G. Kidwell, and P. M. Bingham. 1982. The molecular basis of P-M hybrid dysgenesis: The nature of induced mutations. Cell, 29: 987–994. doi: 10.1016/0092-8674(82)90462-7.

Saito, K. 2013. The epigenetic regulation of transposable elements by PIWI-interacting RNAs in *Drosophila*. Genes & genetic systems, 88: 9–17.

Sambrook, J., and D. W. Russell. 2006. Purification of RNA from Cells and Tissues by Acid Phenol-Guanidinium Thiocyanate-Chloroform Extraction. Cold Spring Harb. Protoc., doi: 10.1101/pdb.prot4045.

Sasa, M. M., P. T. Chippindale, and N. A. Johnson. 1998. Patterns of Postzygotic Isolation in Frogs. Evolution. 52: 1811–1820. doi: 10.2307/2411351.

Schaefer, R. E., M. G. Kidwell, and A. Fausto-Sterling. 1979. Hybrid dysgenesis in Drosophila melanogaster: Morphological and cytological studies of ovarian dysgenesis. Genetics 92: 1141–1152.

Schrider, D. R., J. Ayroles, D. R. Matute, and A. D. Kern. 2018. Supervised machine learning reveals introgressed loci in the genomes of *Drosophila simulans* and *D. sechellia*. PLOS Genet., 14: e1007341. doi: 10.1371/journal.pgen.1007341.

Serrato-Capuchina, A., and D. Matute. 2018. The role of transposable elements in speciation. Genes (Basel). 9: 254.

Serrato-Capuchina, A., J. Wang, E. Earley, D. Peede, K. Isbell, and D. R. Matute. 2020. Paternally inherited *P*-element copy number affects the magnitude of hybrid dysgenesis in *Drosophila simulans* and *D. melanogaster*. Genome Biol. Evol., 12: 808–826, doi: 10.1093/gbe/evaa084.

Shan, X., Z. Liu, Z. Dong, Y. Wang, Y. Chen, X. Lin, L. Long, F. Han, Y. Dong, and B. Liu. 2005. Mobilization of the active MITE transposons *mPing* and *Pong* in rice by introgression from wild rice (*Zizania latifolia* Griseb.). Mol. Biol. Evol. 22: 976–990.

Silva, J. C., and M. G. Kidwell. 2000. Horizontal transfer and selection in the evolution of *P* elements. Mol. Biol. Evol. 17:1542–1557.

Simkin, A., A. Wong, Y. P. Poh, W. E. Theurkauf, and J. D. Jensen. 2013. Recurrent and recent selective sweeps in the piRNA pathway. Evolution 67: 1081–1090. doi: 10.1111/evo.12011.

Siomi, M. C., K. Sato, D. Pezic, and A. A. Aravin. 2011. PIWI-interacting small RNAs: the vanguard of genome defence. Nat. Rev. Mol. Cell Biol. 12: 246–258.

Slotkin, R. K., and R. Martienssen. 2007. Transposable elements and the epigenetic regulation of the genome. Nature Reviews Genetics, 8: 272–285.

Sobel, J. M., G. F. Chen, L. R. Watt, and D. W. Schemske. 2010. The biology of speciation. Evolution 64: 295–315.

Srivastav, S. P., and E. S. Kelleher. 2017. Paternal induction of hybrid dysgenesis in *Drosophila melanogaster* is weakly correlated with both *P*-element and *hobo* element dosage. G3; Genes|Genomes|Genetics, 7: 1487–1497. doi: 10.1534/g3.117.040634.

Srivastav, S. P., R. Rahman, Q. Ma, and N. C. Lau. 2019. *Har-P*, a Short *P*-Element Variant, weaponizes *P*-Transposase to severely impact *Drosophila* gonad development. eLife. 8: e49948.

Dobzhansky. 1937. Genetics and the Origin of Species. Columbia University Press. New York, NY, USA.

Trizzino, M., Y. S. Park, M. Holsbach-Beltrame, K. Aracena, K. Mika, M. Caliskan, G. H. Perry, V. J. Lynch, and C. D. Brown. 2017. Transposable elements are the primary source of novelty in primate gene regulation. Genome Res. 27:1623–1633.

True, J. R., and E. S. Haag. 2001. Developmental system drift and flexibility in evolutionary trajectories. Evolution & Development, 3: 109–119.

Turissini, D. A., A. A. Comeault, G. Liu, Y. C. G. Lee, and D. R. Matute. 2017. The ability of *Drosophila* hybrids to locate food declines with parental divergence. Evolution 71: 960–973.

Turissini, D. A., J. A. McGirr, S. S. Patel, J. R. David, and D. R. Matute. 2018. The rate of evolution of postmating-prezygotic reproductive isolation in *Drosophila*. Mol. Biol. Evol. 35: 312–334.

Ungerer, M. C., S. C. Strakosh, and Y. Zhen. 2006. Genome expansion in three hybrid sunflower species is associated with retrotransposon proliferation. Curr. Biol., 16: R872–R873. doi: 10.1016/j.cub.2006.09.020.

Vaser, R., I. Sović, N. Nagarajan, and M. Šikić. 2017. Fast and accurate de novo genome assembly from long uncorrected reads. Genome Res., 27: 737–746. doi: 10.1101/gr.214270.116.

Vela, D., A. Fontdevila, C. Vieira, and M. Pilar García Guerreiro. 2014. A genome-wide survey of genetic instability by transposition in *Drosophila* hybrids. PLoS One, 9: e88992. doi: 10.1371/journal.pone.0088992.

Venables, W. N. (William N.., B. D. Ripley, and W. N. (William N.). Venables. 2003. Modern Applied Statistics With S. Technometrics 45:111–111.

Vermaak, D., S. Henikoff, and H. S. Malik. 2005. Positive selection drives the evolution of rhino, a member of the heterochromatin protein 1 family in *Drosophila*. PLoS Genet., 1: e9. doi: 10.1371/journal.pgen.0010009.

Wang, L., D. A. Barbash, and E. S. Kelleher. 2020. Adaptive evolution among cytoplasmic piRNA proteins leads to decreased genomic auto-immunity. PLoS Genet., 16: p.e1008861. doi: 10.1371/journal.pgen.1008861.

Warnes, G. R. 2012. gplots: Various R programming tools for plotting data. J. Phycol., doi: 10.1111/j.0022-3646.1997.00569.x.

Warnes, G. R., B. Bolker, L. Bonebakker, R. Gentleman, W. H. A. Liaw, T. Lumley, M. Maechler, A. Magnusson, S. Moeller, M. Schwartz, and B. Venables. 2016. Package “gplots”: Various R programming tools for plotting data. R Packag. version 2.17.0., doi: 10.1111/j.0022-3646.1997.00569.x.

Watson, E. T., and J. P. Demuth. 2013. The role of transposable elements in speciation. Chapter 10: pp. 227–250. Speciation: Natural Processes, Genetics and Biodiversity. Edited by Michalak, P. Nova Science Publishers, Inc. 274 pages

Werren, J. H. 2011. Selfish genetic elements, genetic conflict, and evolutionary innovation. Proc. Natl. Acad. Sci. U. S. A. 108: 10863–10870.

Wong, L. C., and P. Schedl. 2006. Dissection of *Drosophila* ovaries. J. Vis. Exp. 8:52.

