## Supplementary material for "*P*-elements strengthen reproductive isolation within the *Drosophila simulans* species complex": Tables S1-S5, Figure S1-S6

### TABLES S1-S5, FIGURES S1-S6

Antonio Serrato-Capuchina[1,2], Emmanuel R. R. D'Agostino[1], David Peede[1], Baylee Roy[1], Kristin Isbell[1], Jeremy Wang[3], and Daniel R. Matute[1]

[1]Biology Department, University of North Carolina, Chapel Hill, N.C., USA
[2]Department of Organismic and Evolutionary Biology, Harvard University, Cambridge, M.A., USA
[3]Department of Genetics, University of North Carolina, Chapel Hill, N.C., USA

**TABLE S1. SRA numbers of all the reads used in this study.**

| sample | Species | Accession number | Reference |
|---|---|---|---|
| 14021-0428.25 | *sech* | SRS430100 | McManus et al. 2014 |
| Anro_B1 | *sech* | SRS2378192 | Turissini et al. 2018; Schrider et al. 2018 |
| Anro_B2 | *sech* | SRS2378191 | Turissini et al. 2018; Schrider et al. 2018 |
| Anro_B3 | *sech* | SRS2378187 | Turissini et al. 2018; Schrider et al. 2018 |
| Anro_B5 | *sech* | SRS2378177 | Turissini et al. 2018; Schrider et al. 2018 |
| Anro_B6 | *sech* | SRS2378189 | Turissini et al. 2018; Schrider et al. 2018 |
| Anro_B7 | *sech* | SRS2378207 | Turissini et al. 2018; Schrider et al. 2018 |
| Anro_B8 | *sech* | SRS2378279 | Turissini et al. 2018; Schrider et al. 2018 |
| Denis124 | *sech* | SRS2378225 | Turissini et al. 2018; Schrider et al. 2018 |
| Denis135_sampleo5 | *sech* | SRS2378180 | Turissini et al. 2018; Schrider et al. 2018 |
| Denis7_2 | *sech* | SRS2378220 | Turissini et al. 2018; Schrider et al. 2018 |
| DenisAMT_sampleo22 | *sech* | SRS2378209 | Turissini et al. 2018; Schrider et al. 2018 |
| DenisAT3 | *sech* | SRS2378223 | Turissini et al. 2018; Schrider et al. 2018 |
| DenisDNJ6 | *sech* | SRS2378224 | Turissini et al. 2018; Schrider et al. 2018 |
| DenisJT1_sampleo13 | *sech* | SRS2378222 | Turissini et al. 2018; Schrider et al. 2018 |
| DenisMCL_sampleo15 | *sech* | SRS2378182 | Turissini et al. 2018; Schrider et al. 2018 |
| DenisNF100 | *sech* | SRS2378213 | Turissini et al. 2018; Schrider et al. 2018 |
| DenisNF123_1_sampleo3 | *sech* | SRS2378212 | Turissini et al. 2018; Schrider et al. 2018 |
| DenisNF13 | *sech* | SRS2378219 | Turissini et al. 2018; Schrider et al. 2018 |
| DenisNF134 | *sech* | SRS2378208 | Turissini et al. 2018; Schrider et al. 2018 |
| DenisNF155_sampleo14 | *sech* | SRS2378226 | Turissini et al. 2018; Schrider et al. 2018 |
| DenisNF66_sampleo20 | *sech* | SRS2378218 | Turissini et al. 2018; Schrider et al. 2018 |

| | | | |
|---|---|---|---|
| DenisNoni10 | *sech* | SRS2378232 | Turissini et al. 2018; Schrider et al. 2018 |
| DenisNoni101 | *sech* | SRS2378184 | Turissini et al. 2018; Schrider et al. 2018 |
| DenisNoni60 | *sech* | SRS2378221 | Turissini et al. 2018; Schrider et al. 2018 |
| LD11 | *sech* | SRS2378188 | Turissini et al. 2018; Schrider et al. 2018 |
| LD12 | *sech* | SRS2378190 | Turissini et al. 2018; Schrider et al. 2018 |
| LD13 | *sech* | SRS2378195 | Turissini et al. 2018; Schrider et al. 2018 |
| LD14 | *sech* | SRS2378194 | Turissini et al. 2018; Schrider et al. 2018 |
| LD15 | *sech* | SRS2378267 | Turissini et al. 2018; Schrider et al. 2018 |
| LD16 | *sech* | SRS2378196 | Turissini et al. 2018; Schrider et al. 2018 |
| LD8 | *sech* | SRS2378270 | Turissini et al. 2018; Schrider et al. 2018 |
| maria_3 | *sech* | SRS2378281 | Turissini et al. 2018; Schrider et al. 2018 |
| mariane_1 | *sech* | SRS2378193 | Turissini et al. 2018; Schrider et al. 2018 |
| PNF10 | *sech* | SRS2378186 | Turissini et al. 2018; Schrider et al. 2018 |
| PNF11 | *sech* | SRS2378185 | Turissini et al. 2018; Schrider et al. 2018 |
| PNF3 | *sech* | SRS2378178 | Turissini et al. 2018; Schrider et al. 2018 |
| PNF4 | *sech* | SRS2378269 | Turissini et al. 2018; Schrider et al. 2018 |
| PNF5 | *sech* | SRS2378179 | Turissini et al. 2018; Schrider et al. 2018 |
| PNF7 | *sech* | SRS2378176 | Turissini et al. 2018; Schrider et al. 2018 |
| Anro71 | *sech* | TBD | This report |
| Denis72 | *sech* | TBD | This report |
| mau12w | *mau* | SRS300691 | Garrigan et al. 2012; Brand et al. 2013 |
| MauKiti | *mau* | TBD | |
| mauST | *mau* | SRS3302928 | Garrigan et al. 2012; Brand et al. 2013 |
| MS17 | *mau* | SRA058420 | Garrigan et al. 2012; Brand et al. 2013 |

| | | | |
|---|---|---|---|
| R23 | *mau* | SRS691146 | Garrigan et al. 2012; Brand et al. 2013 |
| R31 | *mau* | SRS691151 | Garrigan et al. 2012; Brand et al. 2013 |
| R32 | *mau* | SRS691153 | Garrigan et al. 2012; Brand et al. 2013 |
| R39 | *mau* | SRS691158 | Garrigan et al. 2012; Brand et al. 2013 |
| R41 | *mau* | SRS691179 | Garrigan et al. 2012; Brand et al. 2013 |
| R44 | *mau* | SRS691180 | Garrigan et al. 2012; Brand et al. 2013 |
| R56 | *mau* | SRS691182 | Garrigan et al. 2012; Brand et al. 2013 |
| R61 | *mau* | SRS691271 | Garrigan et al. 2012; Brand et al. 2013 |
| R8 | *mau* | SRS691273 | Garrigan et al. 2012; Brand et al. 2013 |
| R42 | *mau* | TBD | This study |
| R50 | *mau* | TBD | This study |
| Bioko_cascade_1 | *sim* | SAMN13243070 | Matute et al. 2020 |
| Bioko_H1 | *sim* | SAMN13243071 | Matute et al. 2020 |
| Bioko_H9 | *sim* | TBD | Matute et al. 2020 |
| Bioko_LB1 | *sim* | SAMN13243072 | Matute et al. 2020 |
| Bioko_Riaba_9 | *sim* | SAMN13243068 | Matute et al. 2020 |
| Bioko_Riaba_mixed | *sim* | SAMN13243069 | Matute et al. 2020 |
| Kib32 | *sim* | SRX497551 | Rogers et al. 2013 |
| MD06 | *sim* | SRX497558 | Rogers et al. 2013 |
| MD105 | *sim* | SRX497564 | Rogers et al. 2013 |
| MD106 | *sim* | SRX497574 | Rogers et al. 2013 |
| MD15 | *sim* | SRX497559 | Rogers et al. 2013 |
| MD199 | *sim* | SRX495510 | Rogers et al. 2013 |
| MD221 | *sim* | SRX495507 | Rogers et al. 2013 |
| MD233 | *sim* | SRX497557 | Rogers et al. 2013 |
| MD251 | *sim* | SRX497553 | Rogers et al. 2013 |
| MD63 | *sim* | SRX497563 | Rogers et al. 2013 |
| MD73 | *sim* | SRX497560 | Rogers et al. 2013 |
| NS05 | *sim* | SRX497572 | Rogers et al. 2013 |
| NS113 | *sim* | SRX497561 | Rogers et al. 2013 |
| NS137 | *sim* | SRX497561 | Rogers et al. 2013 |
| NS33 | *sim* | SRX497575 | Rogers et al. 2013 |

| | | | |
|---|---|---|---|
| NS39 | *sim* | SRX497562 | Rogers et al. 2013 |
| NS40 | *sim* | SRX497556 | Rogers et al. 2013 |
| NS50 | *sim* | SRX497571 | Rogers et al. 2013 |
| NS67 | *sim* | SRX497565 | Rogers et al. 2013 |
| NS78 | *sim* | SRX497573 | Rogers et al. 2013 |
| NS79 | *sim* | SRX497576 | Rogers et al. 2013 |
| tsimbazazaa | *sim* | SRS430098 | Rogers et al. 2013 |
| w501 | *sim* | SRS1090170 | Rogers et al. 2013 |
| 18BOMA01 | *sim* | TBD | Serrato-Capuchina et al. 2020 |
| 18BOMA03 | *sim* | TBD | Serrato-Capuchina et al. 2020 |
| 18CHAALES15 | *sim* | TBD | Serrato-Capuchina et al. 2020 |
| 18KARI01 | *sim* | TBD | Serrato-Capuchina et al. 2020 |
| 18KARI03 | *sim* | TBD | Serrato-Capuchina et al. 2020 |
| 18KARI04 | *sim* | TBD | Serrato-Capuchina et al. 2020 |
| 18KARI05 | *sim* | TBD | Serrato-Capuchina et al. 2020 |
| 18KARI06 | *sim* | TBD | Serrato-Capuchina et al. 2020 |
| 18KARI09 | *sim* | TBD | Serrato-Capuchina et al. 2020 |
| 18KARI10 | *sim* | TBD | Serrato-Capuchina et al. 2020 |
| 18KARI26 | *sim* | TBD | Serrato-Capuchina et al. 2020 |
| 18KARI37 | *sim* | TBD | Serrato-Capuchina et al. 2020 |
| 18KURITANA20 | *sim* | TBD | Serrato-Capuchina et al. 2020 |
| 18MPALA02C | *sim* | TBD | Serrato-Capuchina et al. 2020 |
| 18MPALA03 | *sim* | TBD | Serrato-Capuchina et al. 2020 |
| 18MPALA05 | *sim* | TBD | Serrato-Capuchina et al. 2020 |
| 18MPALA08 | *sim* | TBD | Serrato-Capuchina et al. 2020 |
| 18MPALA09 | *sim* | TBD | Serrato-Capuchina et al. 2020 |

| | | | |
|---|---|---|---|
| 18MPALA11 | *sim* | TBD | Serrato-Capuchina et al. 2020 |
| 18MPALA12 | *sim* | TBD | Serrato-Capuchina et al. 2020 |
| 18MPLALA13 | *sim* | TBD | Serrato-Capuchina et al. 2020 |
| 18MU04 | *sim* | TBD | Serrato-Capuchina et al. 2020 |
| 18MU05 | *sim* | TBD | Serrato-Capuchina et al. 2020 |
| 18NANY04 | *sim* | TBD | Serrato-Capuchina et al. 2020 |
| 18NANY10 | *sim* | TBD | Serrato-Capuchina et al. 2020 |
| 18NANY15 | *sim* | TBD | Serrato-Capuchina et al. 2020 |
| 18NANY16 | *sim* | TBD | Serrato-Capuchina et al. 2020 |
| KARI25 | *sim* | TBD | Serrato-Capuchina et al. 2020 |
| dsim_florida_116 | *sim* | TBD | Kofler et al. 2015 |

**TABLE S2. Collection details of the lines used for crosses.**

| Species | Line | Location | Collector/Donor |
|---|---|---|---|
| *simulans* | 18MPALA11 | Kenya | Matute DR |
| *simulans* | cascade1 | Equatorial Guinea | Matute DR |
| *simulans* | 18MU05 | Kenya | Matute DR |
| *simulans* | 18KARI10 | Kenya | Matute DR |
| *simulans* | 18MPALA13 | Kenya | Matute DR |
| *simulans* | 18MPALA02C | Kenya | Matute DR |
| *simulans* | BiokoLB1 | Equatorial Guinea | Matute DR |
| *simulans* | Md06 | Madagascar | David JR |
| *simulans* | Md105 | Madagascar | David JR |
| *simulans* | Md106 | Madagascar | David JR |
| *simulans* | Md199 | Madagascar | David JR |
| *simulans* | NS05 | Kenya | Andolfatto P |
| *simulans* | NS40 | Kenya | Andolfatto P |
| *simulans* | NS113 | Kenya | Andolfatto P |
| *sechellia* | LD11 | Seychelles | Matute DR/Ayroles JF |
| *sechellia* | LD16 | Seychelles | Matute DR/Ayroles JF |
| *sechellia* | Denis72 | Seychelles | Matute DR/Ayroles JF |
| *sechellia* | DenisNF13 | Seychelles | Matute DR/Ayroles JF |
| *sechellia* | DenisNF100 | Seychelles | Matute DR/Ayroles JF |
| *sechellia* | Denis124 | Seychelles | Matute DR/Ayroles JF |
| *sechellia* | Anro71 | Seychelles | Matute DR/Ayroles JF |
| *mauritiana* | R8 | Rodrigues | Womack M |
| *mauritiana* | R23 | Rodrigues | Womack M |
| *mauritiana* | R31 | Rodrigues | Womack M |
| *mauritiana* | R32 | Rodrigues | Womack M |
| *mauritiana* | R61 | Rodrigues | Womack M |
| *mauritiana* | R42 | Rodrigues | Womack M |
| *mauritiana* | R50 | Rodrigues | Womack M |

**TABLE S3. Number of dissected females per cross.** If a cross is not listed in this table N=20.

| Cross | Temperature (ºC) | N |
|---|---|---|
| ♀NS113×♂18KARI10 | 23 | 40 |
| ♀NS113×♂18MPALA02C | 23 | 40 |
| ♀NS113×♂18MPALA11 | 23 | 40 |
| ♀NS113×♂18MPLALA13 | 23 | 40 |
| ♀NS113×♂18MU05 | 23 | 40 |
| ♀NS113×♂BiokoLB1 | 23 | 40 |
| ♀NS113×♂cascade1 | 23 | 40 |
| ♀NS113×♂18KARI10 | 29 | 40 |
| ♀NS113×♂18MPALA02C | 29 | 40 |
| ♀NS113×♂18MPALA11 | 29 | 40 |
| ♀NS113×♂18MPLALA13 | 29 | 40 |
| ♀NS113×♂18MU05 | 29 | 40 |
| ♀NS113×♂BiokoLB1 | 29 | 40 |
| ♀NS113×♂cascade1 | 29 | 40 |

**TABLE S4. Hybrid dysgenesis is rare in crosses between ♀*sim* and males from the other species of the *simulans* species complex regardless of the PE genotype of the *D. simulans* female.** We pooled results for each female genotype across seven male types (each cross with N=20 for a total of 140 dissections per mother type).

| Cross | Temperature (ºC) | mother | ♀*sim* PE genotype | Number of dysgenic F1s | Total dissections |
|---|---|---|---|---|---|
| ♀*sim* × ♂*mau* | 23 | 18KARI10 | P | 0 | 140 |
| ♀*sim* × ♂*mau* | 23 | 18MPALA02C | P | 0 | 140 |
| ♀*sim* × ♂*mau* | 23 | 18MPALA11 | P | 0 | 140 |
| ♀*sim* × ♂*mau* | 23 | 18MPLALA13 | P | 0 | 140 |
| ♀*sim* × ♂*mau* | 23 | 18MU05 | P | 0 | 140 |
| ♀*sim* × ♂*mau* | 23 | BiokoLB1 | P | 0 | 140 |
| ♀*sim* × ♂*mau* | 23 | cascade1 | P | 0 | 140 |
| ♀*sim* × ♂*mau* | 23 | Md06 | M | 0 | 140 |
| ♀*sim* × ♂*mau* | 23 | Md105 | M | 0 | 140 |
| ♀*sim* × ♂*mau* | 23 | Md106 | M | 0 | 140 |
| ♀*sim* × ♂*mau* | 23 | Md199 | M | 0 | 140 |
| ♀*sim* × ♂*mau* | 23 | NS05 | M | 0 | 140 |
| ♀*sim* × ♂*mau* | 23 | NS113 | M | 0 | 140 |
| ♀*sim* × ♂*mau* | 23 | NS40 | M | 0 | 140 |
| ♀*sim* × ♂*mau* | 29 | 18KARI10 | P | 0 | 140 |
| ♀*sim* × ♂*mau* | 29 | 18MPALA02C | P | 0 | 140 |
| ♀*sim* × ♂*mau* | 29 | 18MPALA11 | P | 0 | 140 |
| ♀*sim* × ♂*mau* | 29 | 18MPLALA13 | P | 0 | 140 |

| | | | | | |
|---|---|---|---|---|---|
| ♀***sim*** **×** ♂***mau*** | **29** | 18MU05 | P | 0 | 140 |
| ♀***sim*** **×** ♂***mau*** | **29** | BiokoLB1 | P | 0 | 140 |
| ♀***sim*** **×** ♂***mau*** | **29** | cascade1 | P | 0 | 140 |
| ♀***sim*** **×** ♂***mau*** | **29** | Md06 | M | 0 | 140 |
| ♀***sim*** **×** ♂***mau*** | **29** | Md105 | M | 0 | 140 |
| ♀***sim*** **×** ♂***mau*** | **29** | Md106 | M | 0 | 140 |
| ♀***sim*** **×** ♂***mau*** | **29** | Md199 | M | 2 | 140 |
| ♀***sim*** **×** ♂***mau*** | **29** | NS05 | M | 0 | 140 |
| ♀***sim*** **×** ♂***mau*** | **29** | NS113 | M | 1 | 140 |
| ♀***sim*** **×** ♂***mau*** | **29** | NS40 | M | 1 | 140 |
| ♀***sim*** **×** ♂**sech** | **23** | 18KARI10 | P | 0 | 140 |
| ♀***sim*** **×** ♂**sech** | **23** | 18MPALA02C | P | 0 | 140 |
| ♀***sim*** **×** ♂**sech** | **23** | 18MPALA11 | P | 0 | 140 |
| ♀***sim*** **×** ♂**sech** | **23** | 18MPLALA13 | P | 0 | 140 |
| ♀***sim*** **×** ♂**sech** | **23** | 18MU05 | P | 0 | 140 |
| ♀***sim*** **×** ♂**sech** | **23** | BiokoLB1 | P | 0 | 140 |
| ♀***sim*** **×** ♂**sech** | **23** | cascade1 | P | 0 | 140 |
| ♀***sim*** **×** ♂**sech** | **23** | Md06 | M | 0 | 140 |
| ♀***sim*** **×** ♂**sech** | **23** | Md105 | M | 0 | 140 |
| ♀***sim*** **×** ♂**sech** | **23** | Md106 | M | 0 | 140 |
| ♀***sim*** **×** ♂**sech** | **23** | Md199 | M | 0 | 140 |
| ♀***sim*** **×** ♂**sech** | **23** | NS05 | M | 0 | 140 |
| ♀***sim*** **×** ♂**sech** | **23** | NS113 | M | 0 | 140 |

| | | | | | |
|---|---|---|---|---|---|
| ♀***sim*** **×** ♂**sech** | **23** | NS40 | M | 0 | 140 |
| ♀***sim*** **×** ♂**sech** | **29** | 18KARI10 | P | 0 | 140 |
| ♀***sim*** **×** ♂**sech** | **29** | 18MPALA02C | P | 0 | 140 |
| ♀***sim*** **×** ♂**sech** | **29** | 18MPALA11 | P | 0 | 140 |
| ♀***sim*** **×** ♂**sech** | **29** | 18MPLALA13 | P | 0 | 140 |
| ♀***sim*** **×** ♂**sech** | **29** | 18MU05 | P | 0 | 140 |
| ♀***sim*** **×** ♂**sech** | **29** | BiokoLB1 | P | 13 | 140 |
| ♀***sim*** **×** ♂**sech** | **29** | cascade1 | P | 0 | 140 |
| ♀***sim*** **×** ♂**sech** | **29** | Md06 | M | 10 | 140 |
| ♀***sim*** **×** ♂**sech** | **29** | Md105 | M | 7 | 140 |
| ♀***sim*** **×** ♂**sech** | **29** | Md106 | M | 0 | 140 |
| ♀***sim*** **×** ♂**sech** | **29** | Md199 | M | 0 | 140 |
| ♀***sim*** **×** ♂**sech** | **29** | NS05 | M | 0 | 140 |
| ♀***sim*** **×** ♂**sech** | **29** | NS113 | M | 0 | 140 |
| ♀***sim*** **×** ♂**sech** | **29** | NS40 | M | 0 | 140 |

**TABLE S5. PE copy number in the paternal genome affects the number of functional ovaries in conspecific and heterospecific produced F1 females. The analyses are similar to those shown in Table 2 but instead of inferring PE copy number using per-base-pair coverage, we inferred PE copy number from read coverage (as in Serrato-Capuchina et al. 2020).** Each row shows the results for each cross at one of two temperatures (ºC). The linear models for each combination of cross and temperature were binomial models fitted to compare the effect of paternally inherited PEs on conspecific and hybrid F1 females.

| | | **PE Copies** | | | **Cross** | | | **PE copies × cross** | |
|---|---|---|---|---|---|---|---|---|---|
| **Cross** | **ºC** | **Odds ratio CI** | **$X^2_1$** | **P** | **Odds ratio CI** | **F** | **P** | **LRT, $X^2_1$** | **P** |
| ♀*sech* × ♂*sim* | 23 | [-0.386, -0.282] | 158.15 | $< 1\times 10^{-5}$ | [2.044, 3.015] | 103.10 | $< 1\times 10^{-5}$ | 0.223 | 0.637 |
| ♀*sech* × ♂*sim* | 29 | [-0.376 -0.283] | 192.964 | $< 1\times 10^{-5}$ | [0.882, 1.943] | 26.939 | $< 1\times 10^{-5}$ | 9.73 | $1.813 \times 10^{-3}$ |
| ♀*mau* × ♂*sim* | 23 | [-0.383, -0.269] | 125.744 | $< 1\times 10^{-5}$ | [1.712, 2.942] | 59.288 | $< 1\times 10^{-5}$ | 2.447 | 0.118 |
| ♀*mau* × ♂*sim* | 29 | [-0.465, -0.384] | 422.455 | $< 1\times 10^{-5}$ | [0.328, 0.860] | 21.276 | $< 1\times 10^{-5}$ | 0.0447 | 0.833 |

**FIGURE S1.** LD2 for 25% cutoff. Absolute value of the LD2 relative loadings by each of the 27 included TEs. The ranking of TEs in the *x*-axis is the same as in Figure 1B.

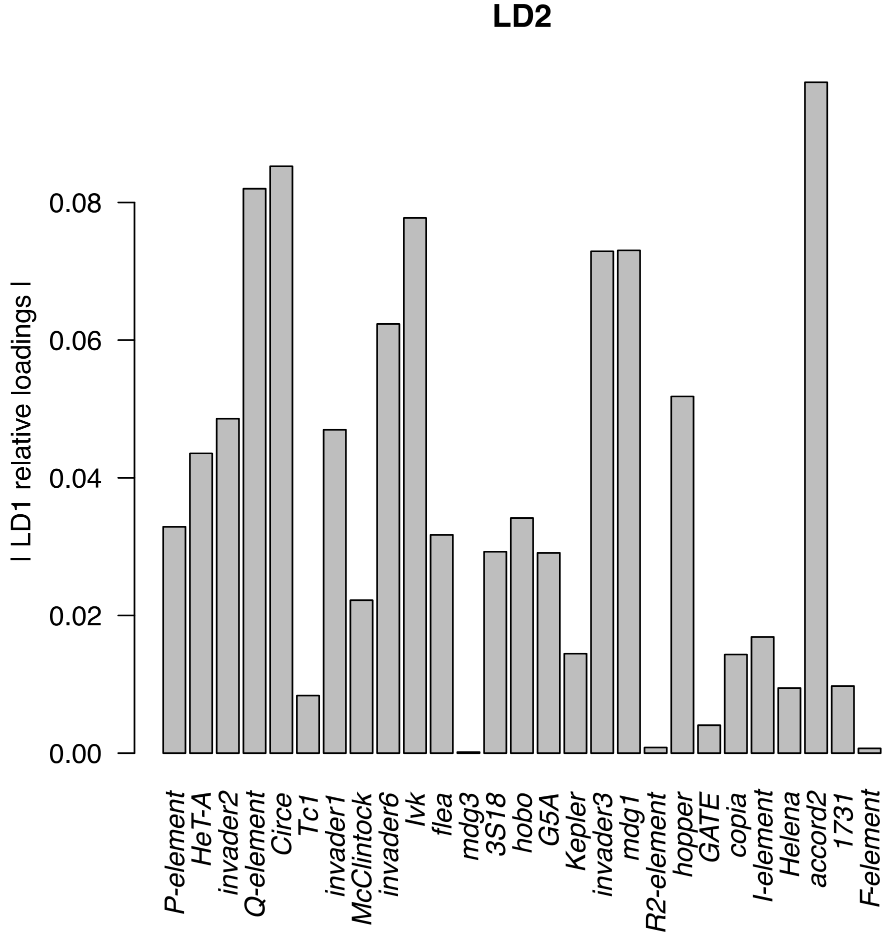

**FIGURE S2. Linear discriminant analysis (LDA) of TEs present in the three species of the *simulans* species complex (0% of missing sequence cutoff).** *D. sechellia*: brown, *D. mauritiana*: purple, *D. simulans*: orange. The axes show the proportion of trace (i.e., separation achieved) by each of the two discriminant functions. Arrows show the 10 TEs with the largest loadings.

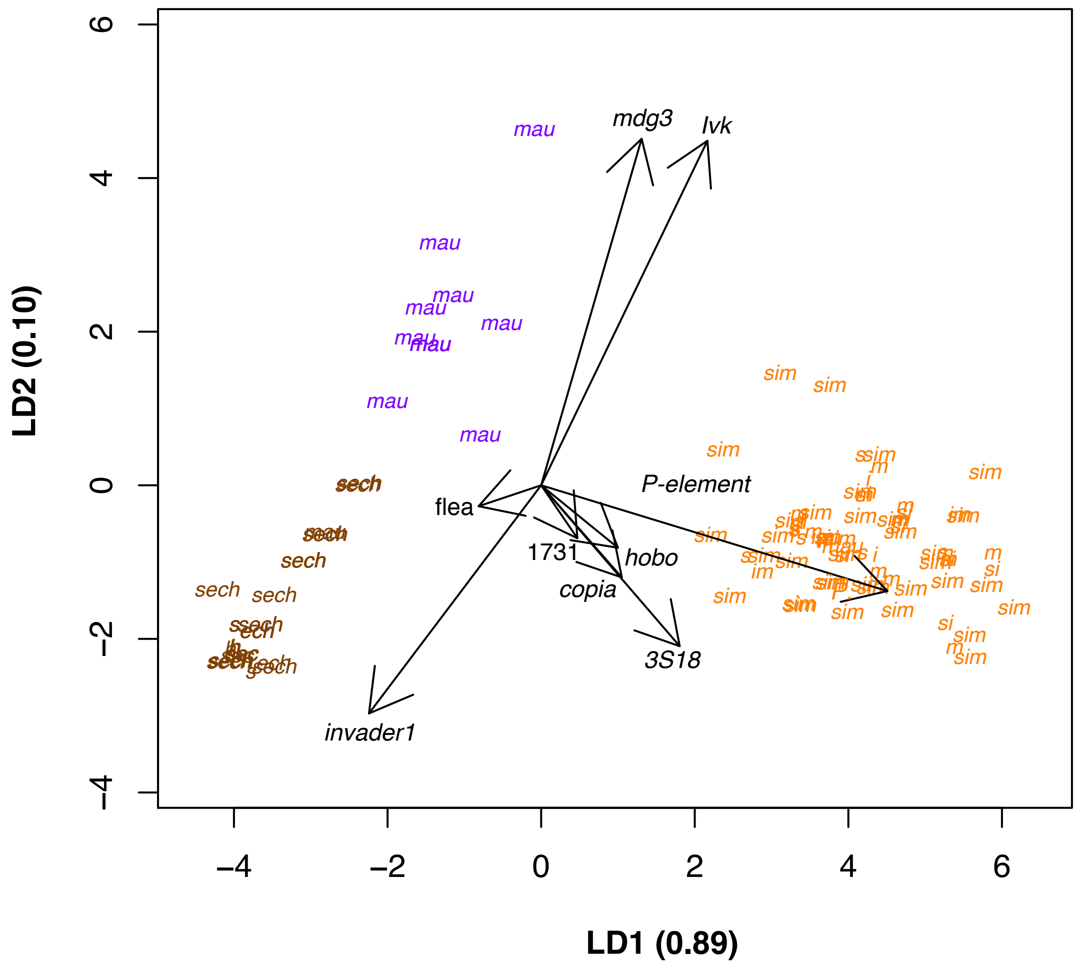

**FIGURE S3. Absolute value of the LD1 and LD2 relative loadings by each of the 15 included TEs when a 0% of missing sequence cutoff is used. A**. LD1. **B**. LD2. The ordering of TEs in the two panels is identical and follows the ranking of relative importance on LD1.

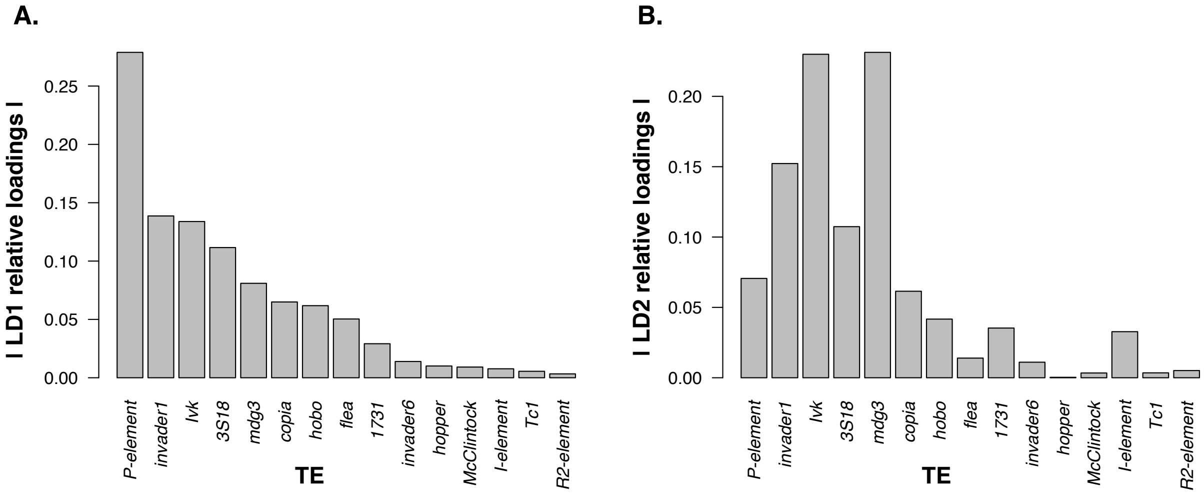

**FIGURE S4. Mean number of copies for different TEs (20) in the *simulans* species complex.** We filtered out copies in which at least 25% of the sequence of the TE was missing. Figure 2 and S5 show the mean numbers for other TEs using the same filtering approach.

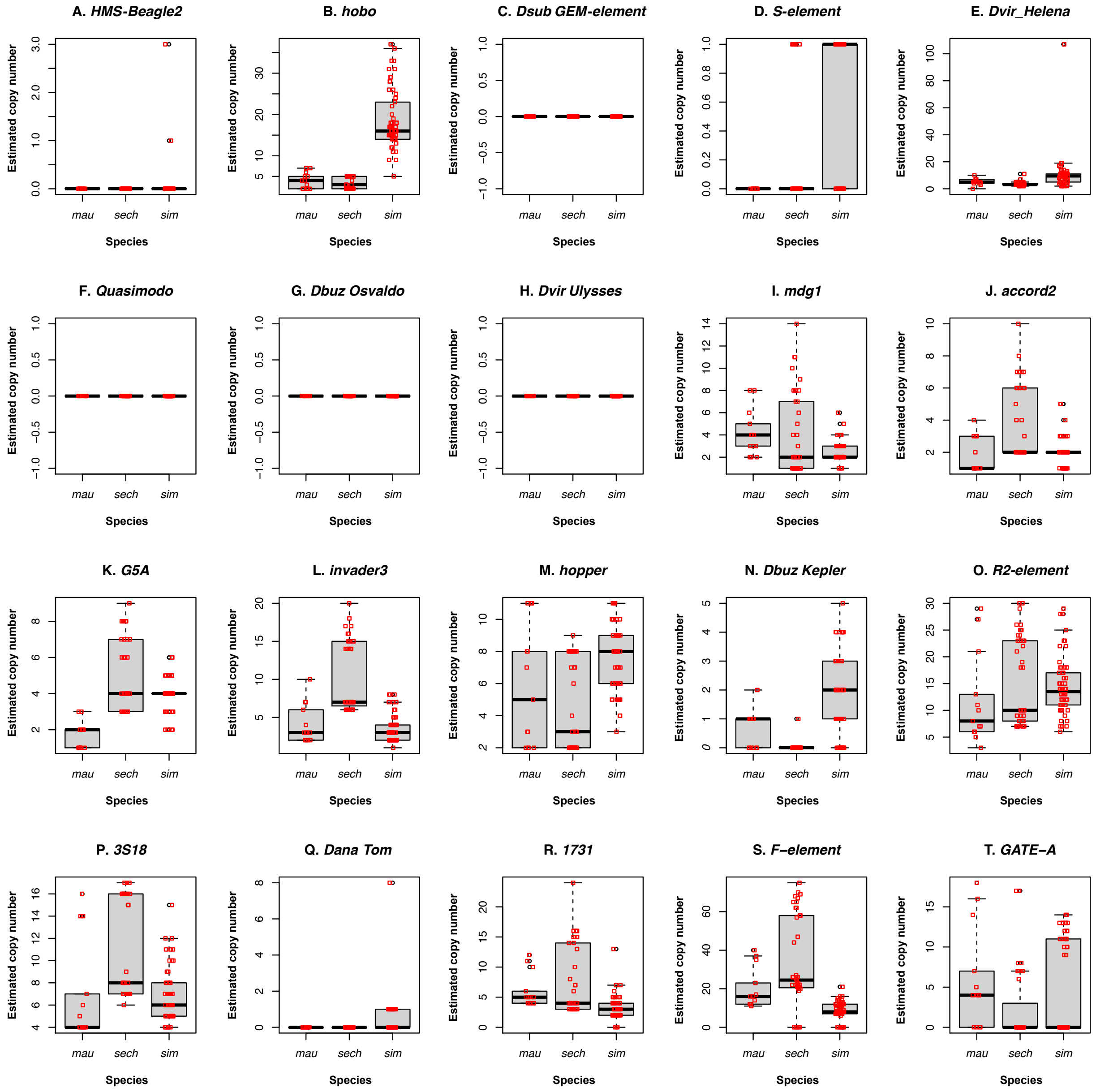

**FIGURE S5**: **Mean number of copies for different TEs (18) in the *simulans* species complex.** We filtered out copies in which at least 25% of the sequence of the TE was missing. Figure 2 and S4 show the mean numbers for other TEs using the same filtering approach.

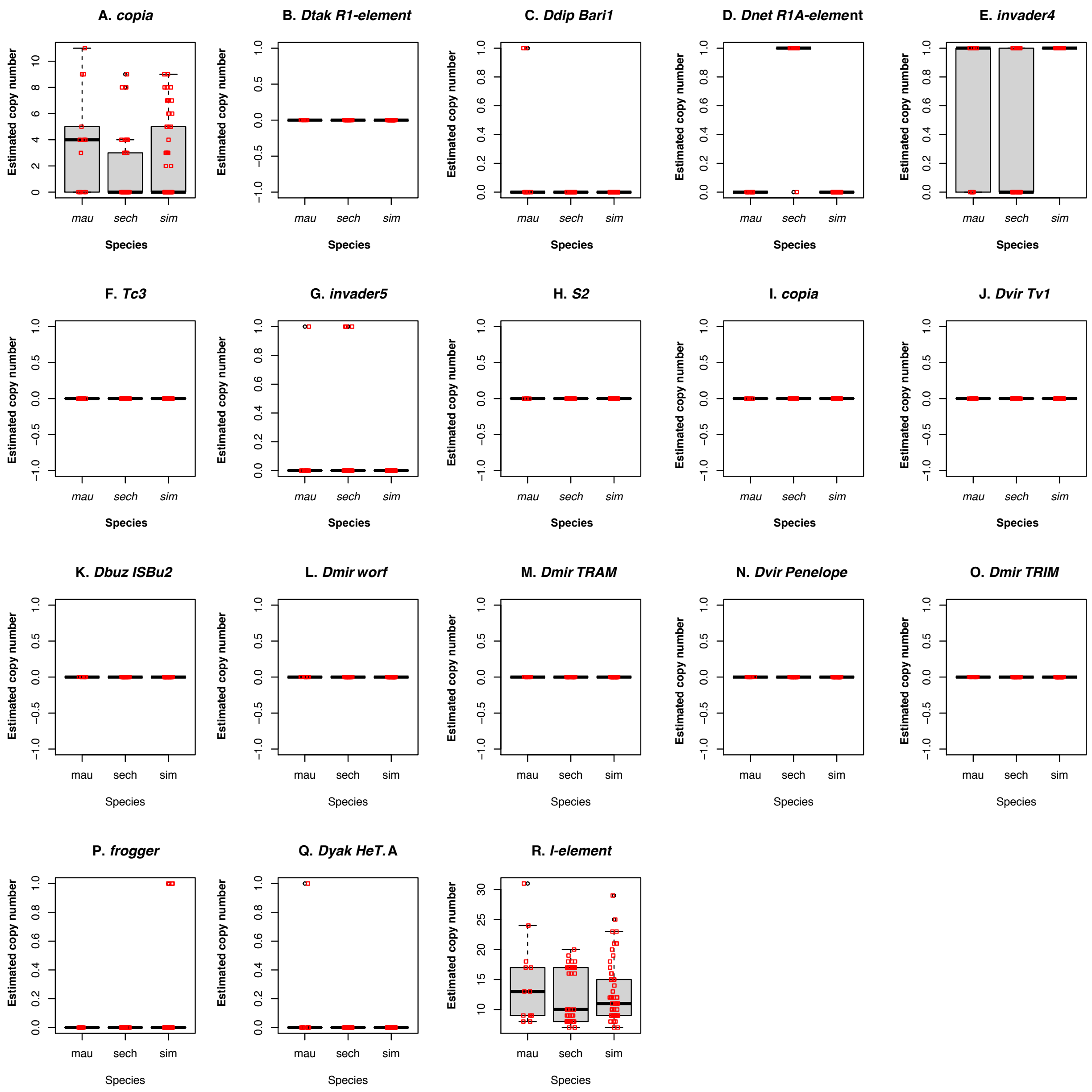

**FIGURE S6**: **Mean number of copies for different TEs (15) in the *simulans* species complex (0%).** The 15 shown TEs had the largest loadings in LD1 (Figure S4A). We filtered out copies in which any the sequence of the TE was missing.

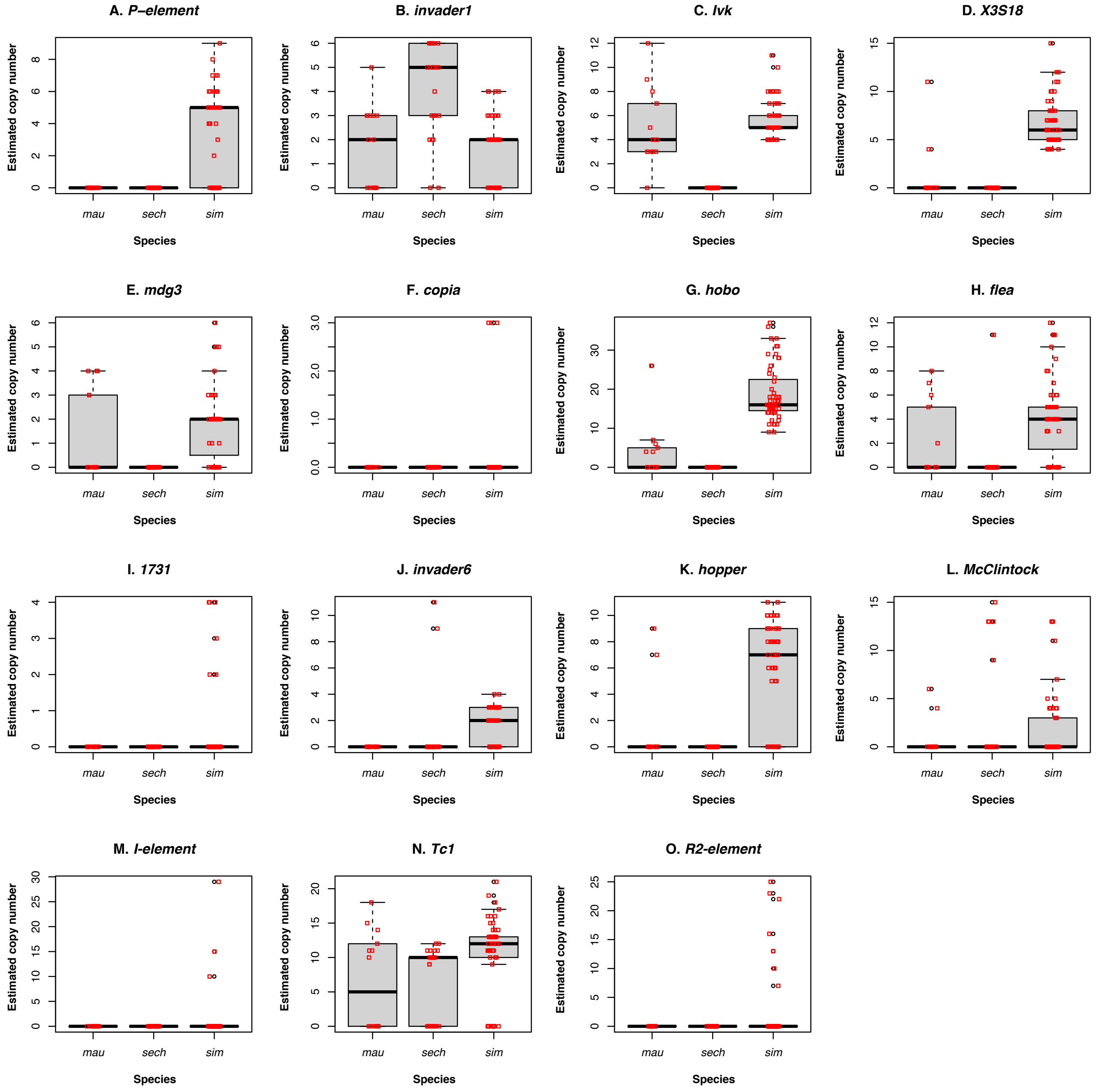
